# Human ORC/MCM density is low in active genes and correlates with replication time but does not solely define replication initiation zones

**DOI:** 10.1101/2020.08.15.252221

**Authors:** Nina Kirstein, Alexander Buschle, Xia Wu, Stefan Krebs, Helmut Blum, Wolfgang Hammerschmidt, Laurent Lacroix, Olivier Hyrien, Benjamin Audit, Aloys Schepers

**Affiliations:** Research Unit Gene Vectors, Helmholtz Zentrum München (GmbH), German Research Center for Environmental Health Marchioninistraße 25, 81377 Munich, Germany; N.K.: current address: University of Miami, Miller School of Medicine, Sylvester Comprehensive Cancer Center, Department of Human Genetics, Biomedical Research Building, 1501 NW 10th Avenue, Miami, FL 33136, USA; Research Unit Gene Vectors, Helmholtz Zentrum München (GmbH), German Research Center for Environmental Health and German Center for Infection Research (DZIF), Partner site Munich, Germany; Institut de Biologie de l’ENS (IBENS), Département de Biologie, Ecole Normale, Supérieure, CNRS, Inserm, PSL Research University, F-75005 Paris, France; X.W. current address: Zhongshan School of Medicine, Sun Yat-sen University, 74 Zhongshan Er Road, Guangzhou, Guangdong Province, China, 510080.; Laboratory for Functional Genome Analysis (LAFUGA), Gene Center of the Ludwig-Maximilians Universität (LMU) München, Feodor-Lynen-Str. 25, D-81377 Munich, Germany; Université de Lyon, ENS de Lyon, Univ Claude Bernard Lyon 1, CNRS, Laboratoire de Physique, F-69342, Lyon, France; Monoclonal Antibody Core Facility, Helmholtz Zentrum München, German Research Center for Environmental Health, Ingolstädter Landstraße 1, D-85764 Neuherberg, Germany

**Keywords:** ORC, MCM complex, ChIP-seq, DNA replication, OK-seq, replication initiation, replication timing, transcription

## Abstract

Eukaryotic DNA replication initiates during S phase from origins that have been licensed in the preceding G1 phase. Here, we compare ChIP-seq profiles of the licensing factors Orc2, Orc3, Mcm3, and Mcm7 with gene expression, replication timing and fork directionality profiles obtained by RNA-seq, Repli-seq and OK-seq. ORC and MCM are strongly and homogeneously depleted from transcribed genes, enriched at gene promoters, and more abundant in early-than in late-replicating domains. Surprisingly, after controlling these variables, no difference in ORC/MCM density is detected between initiation zones, termination zones, unidirectionally replicating and randomly replicating regions. Therefore, ORC/MCM density correlates with replication timing but does not solely regulate the probability of replication initiation. Interestingly, H4K20me3, a histone modification proposed to facilitate late origin licensing, was enriched in late replicating initiation zones and gene deserts of stochastic replication fork direction. We discuss potential mechanisms that specify when and where replication initiates in human cells.

## INTRODUCTION

DNA replication ensures exact genome inheritance. In human cells, replication initiates from 20,000 – 50,000 replication origins selected from a 5-10 fold excess of potential or “licensed” origins (Moiseeva & Bakkenist, 2018; Papior et al., 2012). Origin licensing, also called pre-replicative complex (pre-RC) formation, occurs in late mitosis and during the G1 phase of the cell cycle, when CDK activity is low. During this step, the origin recognition complex (ORC) binds DNA and together with Cdt1 and Cdc6, loads minichromosome maintenance (MCM) complexes, the core motor of the replicative helicase, as inactive head-to-head double hexamers (MCM-DHs) around double-stranded DNA (Bell & Kaguni, 2013; Evrin et al., 2009; Remus & Diffley, 2009). A single ORC reiteratively loads multiple MCM-DHs, but once MCM-DHs have been assembled, neither ORC, nor Cdc6, nor Cdt1 are required for origin activation (Hyrien, 2016; Powell et al., 2015). During S phase, CDK2 and CDC7 kinase activities in conjunction with other origin firing factors convert some of the MCM-DHs into pairs of active CDC45-MCM-GINS helicases (CMG) that nucleate bidirectional replisome establishment (Douglas et al., 2018; Moiseeva & Bakkenist, 2018). MCM-DHs that do not initiate replication are dislodged from DNA during replication.

In the unicellular *S. cerevisiae*, replication origins are genetically defined by specific DNA sequences (Marahrens & Stillman, 1992). In multicellular organisms, no consensus element for origin activity has been identified and replication initiates from flexible locations. Although mammalian origins fire at different times throughout S phase, neighbouring origins tend to fire at comparable times, which partitions of the genome into ∼5,000 replication timing domains (RTDs), (Rivera-Mulia & Gilbert, 2016a). RTDs replicate in a timely coordinated and reproducible order from early to late in S phase (Pope et al., 2014; Zhao et al., 2017). Different, non-exclusive models exist for this temporal regulation. One model suggests that RTDs are first selected for initiation followed by stochastic origin firing within domains (Boulos et al., 2015; Pope et al., 2014; Rhind & Gilbert, 2013; Rivera-Mulia & Gilbert, 2016b). The cascade or domino model suggests that replication first initiates at the most efficient origins within RTDs and then spreads to less efficient origins (Boos & Ferreira, 2019; Guilbaud et al., 2011). Nuclear processes such as transcription have a major impact on replication profiles (Almeida et al., 2018; Chen et al., 2019; Martin et al., 2011). It is believed that flexible chromatin features including DNA and histone modifications and nucleosome dynamics contribute to origin specification (Cayrou et al., 2015; Prioleau & MacAlpine, 2016; O. K. Smith & Aladjem, 2014). For example, H4K20me3 has been proposed to support the licensing of a subset of late replicating origins in heterochromatin (Brustel et al., 2017).

Different approaches have been developed to characterize mammalian origins, at single-molecule level by optical methods or at cell-population level by sequencing purified replication intermediates (Hulke et al., 2020). Sequencing of short nascent strands (SNS-seq and INI-seq) identified initiation sites with high resolution, correlating with transcriptional start sites (TSSs) and CG-rich regions enriched in G-quadruplex motifs (G4s) and CpG-islands (Besnard et al., 2012; Cayrou et al., 2015; Langley et al., 2016; Prioleau & MacAlpine, 2016). Sequencing of replication-bubble containing restriction fragments (bubble-seq) identified origins enriched with DNase hypersensitive regions and the 5’-end but not the body of active transcription units (Mesner et al., 2013). INI-/SNS- and bubble-seq detected a higher origin density in early RTDs than in mid-to-late RTDs (Besnard et al., 2012; Langley et al., 2016; Mesner et al., 2013). Strand-oriented sequencing of Okazaki fragments (OK-seq) revealed the population-averaged replication fork direction (RFD) allowing the mapping of initiation and termination events (Chen et al., 2019; McGuffee et al., 2013; Petryk et al., 2016; D. J. Smith & Whitehouse, 2012).

Consistently with bubble-seq (Mesner et al., 2013) and single molecule studies (Demczuk et al., 2012; Lebofsky et al., 2006), OK-seq studies of human cells (Petryk et al., 2016; Wu et al., 2018) demonstrated that replication initiates in broad but circumscribed initiation zones (IZs) consisting of multiple inefficient sites. OK-seq detected early-firing IZs precisely flanked on one or both sides by actively transcribed genes and late-firing IZs distantly located from active genes. Early IZs are separated from each other by short termination zones (TZs), which are enriched in transcribed genes. Late IZs are separated by very broad, gene-poor TZs (Chen et al., 2019; Petryk et al., 2016). OK-seq also identified unidirectionally replicating regions (URRs) that sometimes separate IZs from TZs, as well as extended regions of null replication fork directionality (NRRs) in gene deserts of uniformly late replication, consistent with temporally late and spatially random initiation and termination (Petryk et al., 2016). When comparing SNS-, bubble-, and OK-seq data, the highest concordance was observed between IZs detected by OK- and bubble-seq (Petryk et al., 2016). Recently, we found an excellent agreement between IZs determined by OK-seq and by EdUseq-HU, which identified newly synthesized DNA in cells synchronously entering S phase in the presence of EdU and hydroxyurea (Tubbs et al., 2018). Within the EdUseq-HU zones, the most efficient sites were associated with poly(dA:dT) tracts but not any of the GC-rich motifs found by SNS-seq (Tubbs et al., 2018). Repli-seq IZs, which are highly consistent with OK-seq IZs, were also recently identified in high-resolution replication timing (RT) profiles (Zhao et al., 2020).

Importantly, the number of IZs identified by OK-seq (5,000 - 10,000) (Petryk et al., 2016) only account for a fraction of the 20,000 - 50,000 initiation events estimated to take place in each S phase. This and replication kinetic considerations led us to postulate that following efficient initiation at “master initiation zones” (Ma-IZs) identified by OK-seq, replication proceeds by cascade activation of secondary zones, which are too dispersed and inefficient to leave an imprint on population-averaged profiles (Petryk et al., 2016). Consistently, single-molecule studies of yeast genome replication by nanopore-sequencing revealed that 10-20% of initiation events occur dispersedly and away from known, efficient origins, in a manner undetectable by cell population methods (Hennion et al., 2020; Muller et al., 2019).

Chromatin immunoprecipitation followed by sequencing (ChIP-seq) is a complementary method to map ORC and MCM chromatin binding. *Drosophila* ORC ChIP-seq suggests a stochastic binding pattern often correlating with open chromatin marks found at TSSs (MacAlpine et al., 2010). MCM ChIP-seq revealed that MCMs are initially loaded at ORC binding sites in absence of Cyclin E/CDK2 activity, but when Cyclin E/CDK2 activity rises in late G1, MCMs are more abundantly loaded and redistributed, resulting in a loss of spatial overlap with ORC together with MCM-DH displacement from actively transcribed genes (Powell et al., 2015). In human cells, ChIP-seq experiments with single ORC subunits led to the identification of 13,000 to 52,000 potential ORC binding sites (Dellino et al., 2013; Miotto et al., 2016). In a previous study, we used the Epstein-Barr virus (EBV) as model system to compare ORC binding and replication initiation. During latency, replication of the EBV genome is entirely dependent on the human replication initiation machinery. A 5-10-fold excess of potential origins are licensed per genome with respect to 1-3 initiation event(s) mapped (Norio & Schildkraut, 2001, 2004; Papior et al., 2012). A recent genome-wide Mcm7 binding study in human HeLa cells proposed that MCM-DHs bind in excess regardless of the chromatin environment, but that origin activation preferentially occurs upstream of active TSSs (Sugimoto et al., 2018).

Here, we present the first comparative survey of four different pre-RC components and replication initiation events in the human genome by combining ChIP-seq and OK-seq analyses in the lymphoblastoid Raji cell line. We perform ORC and MCM ChIP-seq in cell cycle synchronized pre-replicative (G1) chromatin, to obtain a comprehensive picture of ORC/MCM behavior before replication. We find ORCs and MCMs broadly distributed over the genome, where ORC density highly correlates with early replication timing, an effect less prominently observed for MCMs. We observe that active transcription locally influences ORC and MCM distribution. ORC/MCM are displaced from actively transcribed gene bodies and enriched at active gene promoters. ORC/MCM density is homogeneous over non-transcribed genes and intergenic regions of comparable RT. Consequently, MCM depletion characterizes genic borders of early IZs but not of other IZ borders. Furthermore, URRs, which are refractory to replication initiation, show a similar ORC/MCM density as IZs and TZs. NRRs do not show higher ORC/MCM densities than IZs, TZs and URRs. These findings suggest that ORC/MCM densities do not solely determine IZs and that a specific contribution of the local chromatin environment is required. Indeed, we previously showed that IZs are enriched in open chromatin marks typical of active or poised enhancers (Petryk et al., 2016). Here, we further show that a subset of non-genic late IZs are enriched in H4K20me3, confirming our previous finding that H4K20me3 enhances origin activity in certain chromatin environments (Brustel et al., 2017; Shoaib et al., 2018).

Our findings support the cascade model for replication initiation: although, on average, the entire genome (except transcribed genes) is licensed for replication initiation, adjacent active transcription and internal epigenetic marks specify the extent of Ma-IZs, from which diverging forks emanate before secondary origin activation takes place.

## Results

### Moderate averaging is the best approach for ORC and MCM-DH distribution analysis

To obtain a complete picture of ORC and MCM distributions prior to replication initiation, we cell-cycle fractionated human lymphoblastoid Raji cells by centrifugal elutriation into a pre-replicative G1 population, which is enriched for ORC/MCM bound chromatin (Papior et al., 2012). Propidium Iodide staining followed by FACS (Suppl. Fig. 1a) and Western blot analyses of cyclins A, B, and H3S10 phosphorylation (Suppl. Fig. 1b) confirmed the cell cycle stages. To ensure unbiased detection of ORC and MCM positions by ChIP-seq, we simultaneously targeted two members of both complexes: Orc2, Orc3, Mcm3 and Mcm7, using validated ChIP-grade antibodies (validated in: Papior et al., 2012; Ritzi et al., 2003; Schepers et al., 2001). ChIP efficiency and quality were measured using the Epstein-Barr virus latent origin *oriP* as reference (Suppl. Fig. 1c). The Raji cell line contains 50-60 EBV episomes (Adams et al., 1973). The viral protein EBNA1 recruits ORC to *oriP’s* dyad symmetry element, followed by MCM-DH loading. We detected both ORC and MCM at the dyad symmetry element, as expected (Papior et al., 2012; Ritzi et al., 2003).

ChIP-sequencing of two ORC (Orc2, Orc3) and three MCM (Mcm3, Mcm7) replicates resulted in reproducible, but dispersed ChIP-seq signals as exemplified at the well-characterized replication origin Mcm4/PRKDC (Fig. 1a). We first employed the MACS2 peak-calling program (Feng et al., 2012; Zhang et al., 2008), but found that the obtained results were too dependent on the chosen program settings. At genome-wide levels, we found that ORC and MCM distributions were too dispersed to be efficiently captured by peak calling (data not shown), requiring an alternative approach.

**Figure 1:**
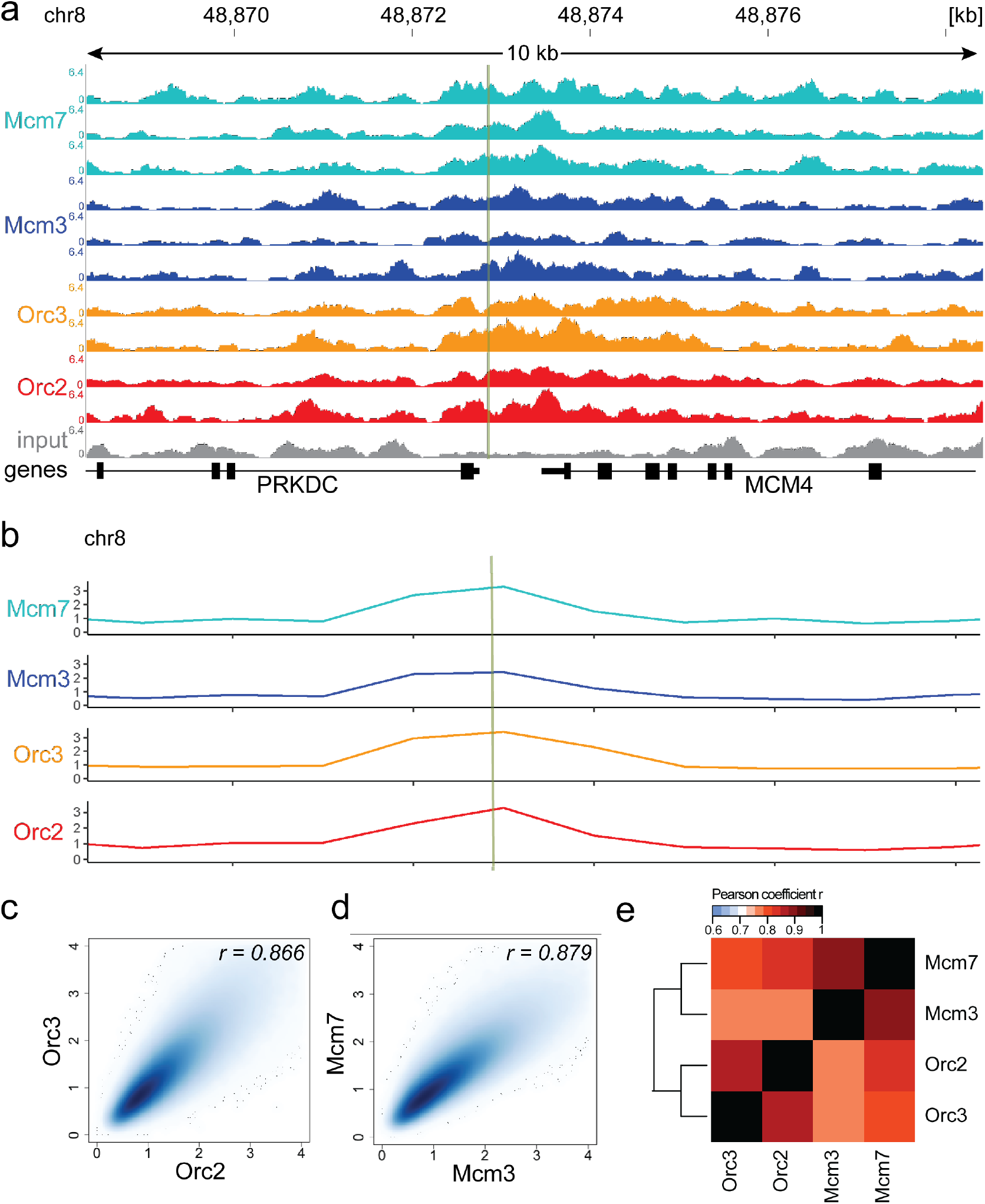
ORC/MCM ChIP-seq is best analyzed using a moderate averaging approach. a) Sequencing profile visualization in UCSC Genome Browser (hg19) at the Mcm4/PRKDC origin after RPGC normalization: two samples of Orc2, Orc3 and three samples of Mcm3, Mcm7, plotted against the input. The profiles are shown in a 10 kb window (chr8: 48,868,314 - 48,878,313), the position of the origin is indicated as green line. b) The profile of ORC/MCM ChIP-seq after 1 kb binning at the same locus. The reads of replicates were summed and normalized by the total genome-wide ChIP read frequency followed by input division. Y-axis represents the resulting relative read frequency. c) Correlation plot between Orc2 and Orc3 relative read frequencies in 1 kb bins. d) Correlation plot between Mcm3 and Mcm7 relative read frequencies in 1 kb bins. e) Heatmap of Pearson correlation coefficients r between all ChIP relative read frequencies in 1 kb bins. Column and line order were determined by complete linkage hierarchical clustering using the correlation distance (d = 1-r).

Consequently, we summed up the reads of the ChIP replicates in 1 kb bins and normalized the signals against the mean read frequencies of each ChIP sample and against input, as is standard in most ChIP-seq analyses. However, in line with a previous report (Teytelman et al., 2009), the input sequencing control was differentially represented in DNase HS regions or regions of biological function. For example, the input was significantly underrepresented in DNase HS regions (DNase HS clusters retrieved from an ENCODE dataset comprising 125 cell lines, see Material and Methods section; Suppl. Fig. 2a), TSSs (Suppl. Fig. 2b), and early RTDs (Suppl. Fig. 2c). As sonication-hypersensitive regions correlate with DNase HS regions (Schwartz et al., 2005), we carefully compared our input-normalized approach versus no input normalization: some quantitative but no qualitative changes were observed. We have included figures showing non-normalized data in the Supplement. Furthermore, the 1 kb bin size was large enough to average out stochastic variations of ORC/MCM, but small enough to detect significant local changes. For example, ORC/MCM enrichments at the Mcm4/PRKDC origin were detected after binning (Fig. 1b, Suppl. Fig. 3a without input division). The relative read frequencies of Orc2/Orc3 (Fig. 1c, Suppl. Fig. 3b without input division) and Mcm3/Mcm7 (Fig. 1d, Suppl. Fig. 3c) showed high Pearson correlation coefficients of r = 0.866 and r = 0.879, respectively. The correlations between ORC and MCM were only slightly lower (Mcm3/Orc2/3: r = 0.775/0.757, Mcm7/Orc2/3: r = 0.821/0.800, Fig. 1e, Suppl. Fig. 3d). Hierarchical clustering based on Pearson correlation distance between ChIP profiles showed that ORC and MCM profiles clustered together. We conclude that this binning approach is valid for analyzing our ChIP-seq data.

Miotto et al. (2016) demonstrated that Orc2 positions highly depend on chromatin accessibility and colocalize with DNase hypersensitive (HS) sites present at active promoters and enhancers. Furthermore, Sugimoto et al. (2018) observed that active origins, enriched with Mcm7 correlate with open chromatin sites. We indeed found a significant enrichment of ORC/ MCM at DNase HS regions larger than 1 kb, compared to regions deprived of DNase HS sites (Suppl. Fig. 3e), which further validated our data.

### ORC/MCM are enriched in initiation zones dependent on transcription

After confirming the validity of the ChIP experiments and establishing an analysis approach based on moderate averaging, we compared the relative read frequencies of ORC/MCM to active replication initiation units. Using OK-seq in Raji cells (Wu et al., 2018), we calculated the RFD (see methods), and delineated zones of preferential replication initiation as ascending segments (AS) of the RFD profile. OK-seq detects RFD upshifts that define origins to kb resolution in yeast (McGuffee et al., 2013), but in mammalian cells these transitions are more gradual, extending over 10-100 kb (Chen et al., 2019; McGuffee et al., 2013; Petryk et al., 2016; Tubbs et al., 2018; Wu et al., 2018). To assess ChIP signals within AS, we only kept ASs > 20 kb. Using the RFD shift across the ASs (ΔRFD) as a measure of replication initiation efficiency, we further required ΔRFD > 0.5 to make sure ASs corresponded to efficient IZs. In total, we selected 2,957 ASs, with an average size of 52.3 kb, which covered 4.9% (155 Mb) of the genome (Fig. 2a, green bars, Table 1). 2,451 (83%) of all AS located close to genic regions (ASs extended by 20 kb on both sides overlapped with at least one annotated gene). 673 ASs (22.8% of all AS) were flanked by actively transcribed genes (TPM > 3; TPM: transcript per million) on both sides (type 1 AS) with less than 20 kb between AS borders and the closest transcribed gene. 1,026 ASs (34.7%) had only one border associated to a transcribed gene with TPM > 3 (type 2 AS). 506 ASs (17.1%) were devoid of proximal genes (non-genic AS), where 20 kb extended ASs did not overlap with any annotated gene (Table 1). Although the slope did not change considerably in the different AS types, type 1 ASs were on average the most efficient, while non-genic ASs were slightly less efficient (Suppl. Fig. 4a). Furthermore, type 1 and type 2 ASs located to early replication timing domains, while non-genic AS were predominantly late replicating (Suppl. Fig. 4b), which is in agreement with IZ previously described for GM06990 and HeLa (Petryk et al., 2016).

**Figure 2:**
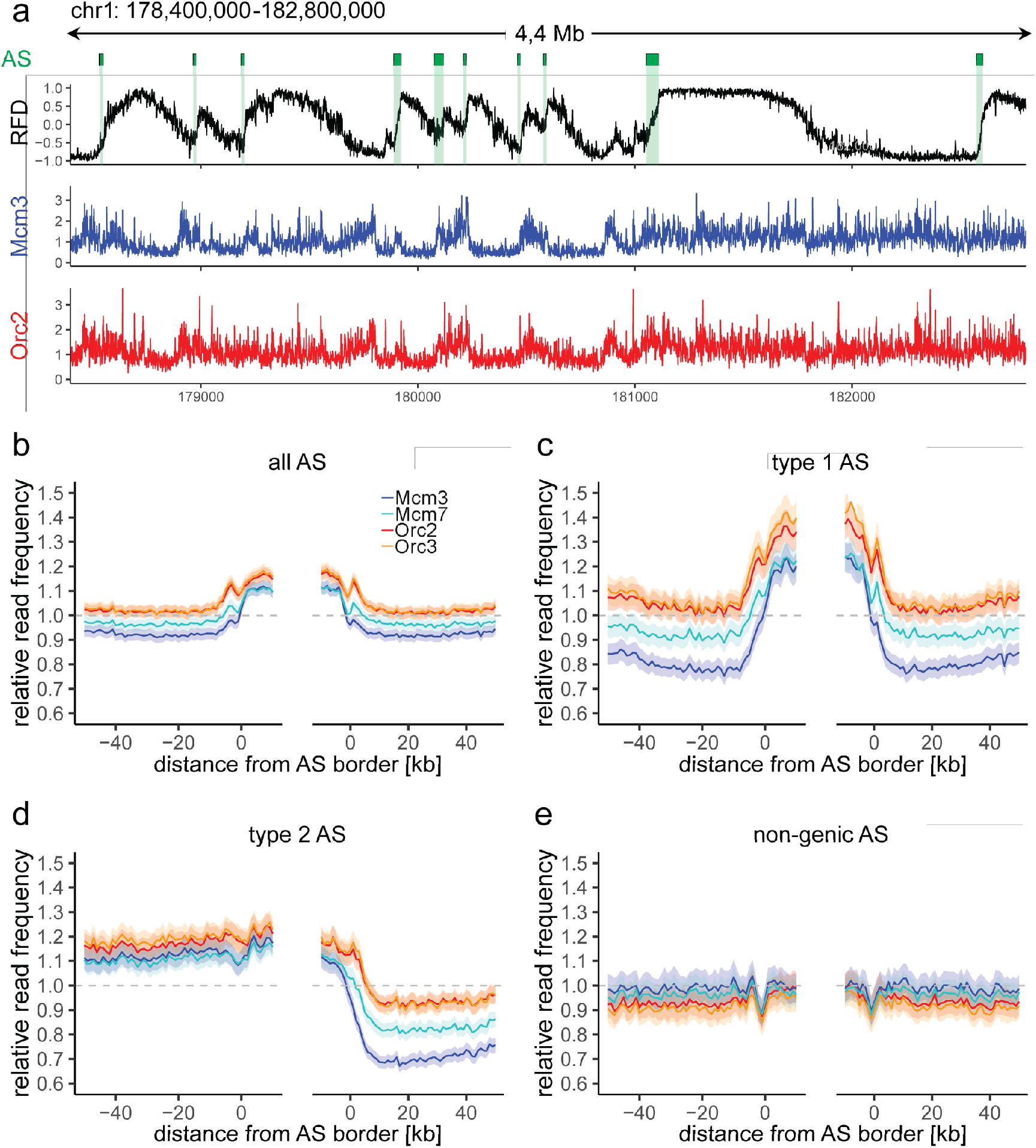
ORC/MCM enrichment within AS depends on active transcription. a) Top panel: Example of an RFD profile on chr1: 178,400,000 – 182,800,000, covering 4 Mb. Detected ASs are labeled by green rectangles. Middle and bottom panel: Representative Mcm3 (blue) and Orc2 (red) ChIP-seq profiles after binning for the same region. b-e) Average input-normalized relative ChIP read frequencies of Orc2, Orc3, Mcm3, and Mcm7 at AS borders of a) all AS (n = 2957), c) type 1 ASs with transcribed genes at both AS borders (n = 673), d) type 2 ASs oriented with their AS border associated to a transcribed genes at the right (n = 1026), and e) non-genic ASs in gene deprived regions (n = 506). The mean of ORC and MCM relative read frequencies are shown ± 2 x SEM (lighter shadows). The dashed grey horizontal line indicates relative read frequency 1.0 for reference.

**Table 1:** Characterization of different AS subtypes. Only AS ≥ 20kb with ΔRFD>0.5 were considered. Genic ASs: ASs extended 20 kb on both sides is overlapped by genic region(s) irrespective of transcriptional activity. Type 1 and type 2 AS: AS flanked by expressed genes (TPM ≥ 3) within 20kb on both sides (type 1) or one side (type 2). Non-genic: no annotated gene ± 20kb of AS border.

|  | Number | Genome coverage [%] | Average length [kb] |
| --- | --- | --- | --- |
| All AS | 2,957 | 4.9 | 52.3 |
| Genic AS | 2,451 | 4.1 | 52.3 |
| Type 1 AS | 673 | 1.1 | 50.7 |
| Type 2 AS | 1,026 | 5.2 | 50.2 |
| Non-genic AS | 506 | 0.8 | 50.7 |

Replication can only be activated when functional pre-RCs are established in the preceding G1 phase. We set our ORC/MCM ChIP-seq signals in relation to RFD and computed the relative read frequencies of ORC/MCM around all AS aggregate borders. Both ORC and MCM were enriched within ASs compared to flanking regions (Fig. 2b, Suppl. Fig. 5a without input division).

To resolve the impact of transcriptional activity, we repeated this calculation and sorted for type 1 ASs (Fig. 2c, Suppl. Fig. 5b), type 2 ASs (Fig. 2d, Suppl. Fig. 5c), and non-genic ASs (Fig. 2e, Suppl. Fig. 5d). Transcriptional activity in AS flanking regions was associated with increased ORC/MCM levels inside ASs (comparing Fig. 2b and Fig. 2c) and a prominent depletion of MCMs from transcribed regions (Fig. 2c and 2d). In contrast, in type 2 ASs, ORC/MCM levels remained elevated at non-transcribed AS borders (Fig. 2d, left border). No evident ORC/MCM enrichments were detected within non-genic ASs (Fig. 2e).

AS borders associated with transcriptional activity were locally enriched in ORC/MCM (Fig. 2c and 2d (right border)). This is in line with previously detected Orc1 accumulation at IZ borders (Petryk et al., 2016). Reciprocally, non-genic AS borders showed a local dip in ORC/MCM levels (Fig. 2d (left border) and 2e), but the biological significance of these observations is unclear. A sequence analysis revealed biased distributions of homopolymeric repeat sequences at AS borders (data not shown). Such sequences may affect nucleosome formation and ORC binding, but may also bias Okazaki fragment detection, hence border detection, at small scales, which explains the local RFD peaks observed at AS borders (Suppl. Fig. 4a).

### ORC and MCM are depleted from transcribed gene bodies and enriched at TSSs

Replication initiation and termination often correlate with active gene transcription (Besnard et al., 2012; Cayrou et al., 2015; Cayrou et al., 2011; Picard et al., 2014). Consistent with previous OK-seq studies (Chen et al., 2019; Petryk et al., 2016), we find on average a strong ascending RFD signal upstream of TSSs and downstream of TTSs of active genes, and a negative RFD slope across the active gene bodies (Suppl. Fig. 6a). This behavior depends on transcriptional activity, as silent genes display an overall flat RFD profile (Suppl. Fig. 6a). The ORC/MCM enrichment in type 1 and 2 ASs compared to flanking genic regions (Fig. 2c and d) argues for a major contribution of active transcription to ORC/MCM positioning. To study this, we set our ChIP-seq data in relation to transcription profiles obtained from asynchronously cycling Raji cells. We analyzed ORC/MCM relative read frequencies around active TSSs and transcriptional termination sites (TTSs) (Fig. 3, Suppl. Fig. 6b-e without input normalization). ORC relative read distribution of G1-phased cells was significantly enriched at active TSSs as already demonstrated in *Drosophila* (MacAlpine et al., 2010) and human cells (Dellino et al., 2013; Miotto et al., 2016). ORC levels were located moderately but significantly higher upstream of TSSs and downstream of TTSs than within active genes (Fig. 3a, Suppl. Fig. 6b). The depletion of ORC from gene bodies was statistically significant for approximately 45% of actively transcribed genes (Suppl. Table 1). Compared to ORC, Mcm3 and Mcm7 enrichments at TSSs were less prominent, but depletion from gene bodies was more pronounced (Fig. 3a, Suppl. Fig. 6b), with 75% and 58% of investigated transcribed gene bodies significantly depleted from Mcm3 and Mcm7, respectively (Suppl. Table 1).

**Figure 3:**
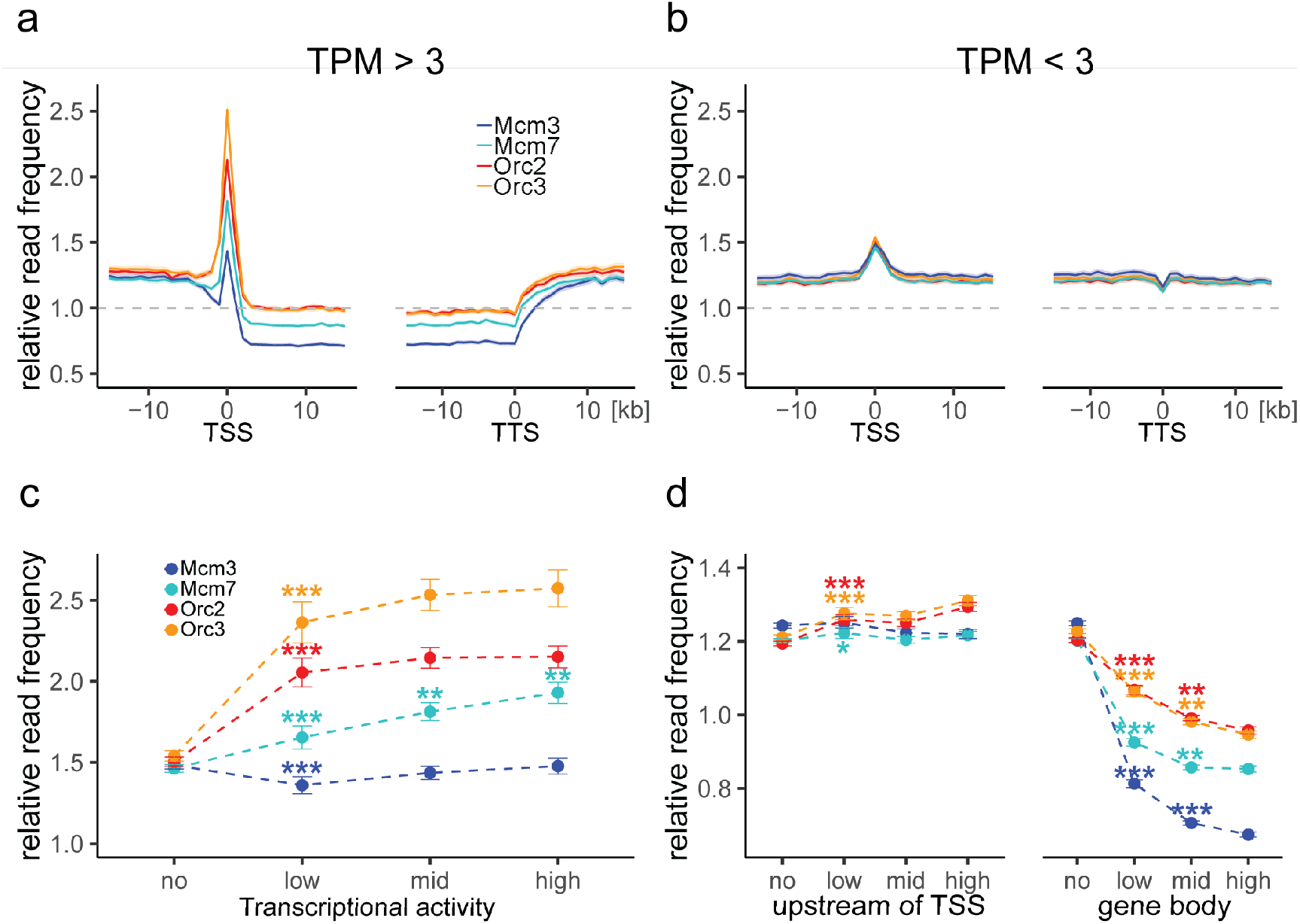
ORC is enriched at active TSSs while MCM is depleted from actively transcribed genes. a) - b) ORC/MCM relative read frequencies around TSSs or TTSs for a) active genes (TPM > 3) and b) inactive genes (TPM < 3). Only genes larger than 30 kb without any adjacent gene within 15 kb were considered. Distances from TSSs or TTSs are indicated in kb. Means of ORC and MCM frequencies are shown ± 2 x SEM (lighter shadows). The dashed grey horizontal line indicates relative read frequency 1.0 for reference. c) ORC/MCM relative read frequencies at TSSs dependent on transcriptional activity (± 2 x SEM). d) ORC/MCM relative read frequencies upstream of TSSs and within the gene body dependent on transcriptional activity (± 2 x SEM; TSSs ± 3 kb removed from analysis). Transcriptional activity was classified as: no (TPM < 3), low (TPM 3-10), mid (TPM 10-40), high (TPM > 40). Statistics were performed by one-way ANOVA followed by Tukey’s post-hoc test. P-values are indicated always comparing to the previous transcriptional level. * p < 0.05, ** p < 0.01, *** p < 0.001.

Depletion was strictly homogeneous from TSS to TTS, strongly suggesting that transcription itself displaces ORC and MCM-DH complexes (Fig. 3a). In contrast, at silent genes, ORC/MCM were hardly enriched at TSSs and were not depleted from gene bodies (Fig. 3b, Suppl. Fig. 6c without input-correction). This confirms that active transcription is required for TSS enrichment and gene body depletion of ORC/MCM. Nevertheless, we also observed that increasing transcriptional activity (classified as follows: low: 3-10 TPM, mid: 10-40 TPM, high: > 40 TPM) did not have any major impact on ORC/MCM enrichments at TSSs (Fig. 3c, Suppl. Fig. 6d without input-correction). ORC/MCM depletion within gene bodies slightly increased with transcription levels (Fig. 3d), but this was less convincing in the non-input-normalized data (Suppl. Fig. 6e). Basal ORC/MCM levels upstream of TSSs and downstream of TTSs were identical, indicating that the local ORC enrichment at TSSs does not result in more MCM loading upstream than downstream of active genes.

Pre-replicative chromatin represents a cell cycle stage immediately prior to replication initiation, with an excess of MCM-DH loaded onto chromatin (Powell et al., 2015; Takahashi et al., 2005). Our observation that Mcm3 and Mcm7 are significantly depleted from gene bodies is consistent with their active displacement by transcription, as previously proposed in *Drosophila* (Powell et al., 2015).

### ORC/MCM genomic distribution is broad and correlated with replication timing but not initiation zones

In the previous sections, we observed a striking influence of transcriptional activity on ORC/MCM distribution that contributes to circumscribing IZs. However, transcriptionally active regions only account for a small subset of the genome. Instead, at non-genic AS, a rather homogenous ORC/MCM pattern within and outside AS is visible (Fig. 2e, Suppl. Fig. 5d without input-correction).

Replication timing (RT) is a crucial aspect of genome stability that has been strongly correlated with gene expression and chromatin structure. RT is determined by the timing of origin firing. In yeast, it has been proposed that the number of MCM-DHs loaded at origins correlates with RT, suggesting how RT profiles can emerge from stochastic origin firing (Das et al., 2015; Yang et al., 2010). In yeast, however, MCM-DHs are only detectable at origins, whereas in human cells ORC/MCM are broadly distributed within and outside IZs so that the presence of ORC/MCM does not solely define RT.

To clarify the relationships between IZ location, IZ firing time and ORC/MCM density in human cells, we used Raji Early/Late Repli-seq data from Sima *et al*. (Sima et al., 2019) and related RT to ORC/MCM relative read frequencies and RFD slope. Thus, we analyzed i) ascending RFD segments (ASs), corresponding to IZs, ii) descending RFD segments (DSs), determined symmetrically to ASs and representing predominant replication termination, iii) unidirectional replicating regions (URRs; segments were |RFD| > 0.8 over > 300 kb), and iv) regions of null RFD i.e. bidirectional replication (NRRs; segments were |RFD| < 0.15 over > 500 kb). Example representations of these four categories are depicted in Suppl. Fig. 7a and 7b. In Fig. 4 we calculated relative Orc2 and Mcm3 (see Suppl. Fig. 7 d-e for Orc3 and Mcm7) read frequencies in 10 kb bins against RT in intergenic regions (left column), silent (TPM < 3) genes (middle column), and active (TPM > 3) genes (right column), while considering either all bins (top row) or bins corresponding to ASs, DSs, URRs, and NRRs (following rows in descending order). ChIP frequencies are normalized by column, i.e. each column is the probability density function of ChIP frequency at a given RT bin. Intergenic regions (left column of each panel) show an ORC/MCM distribution mostly independent from RFD slope, but depending on RT, with higher ORC and MCM levels in early RTDs.

**Figure 4:**
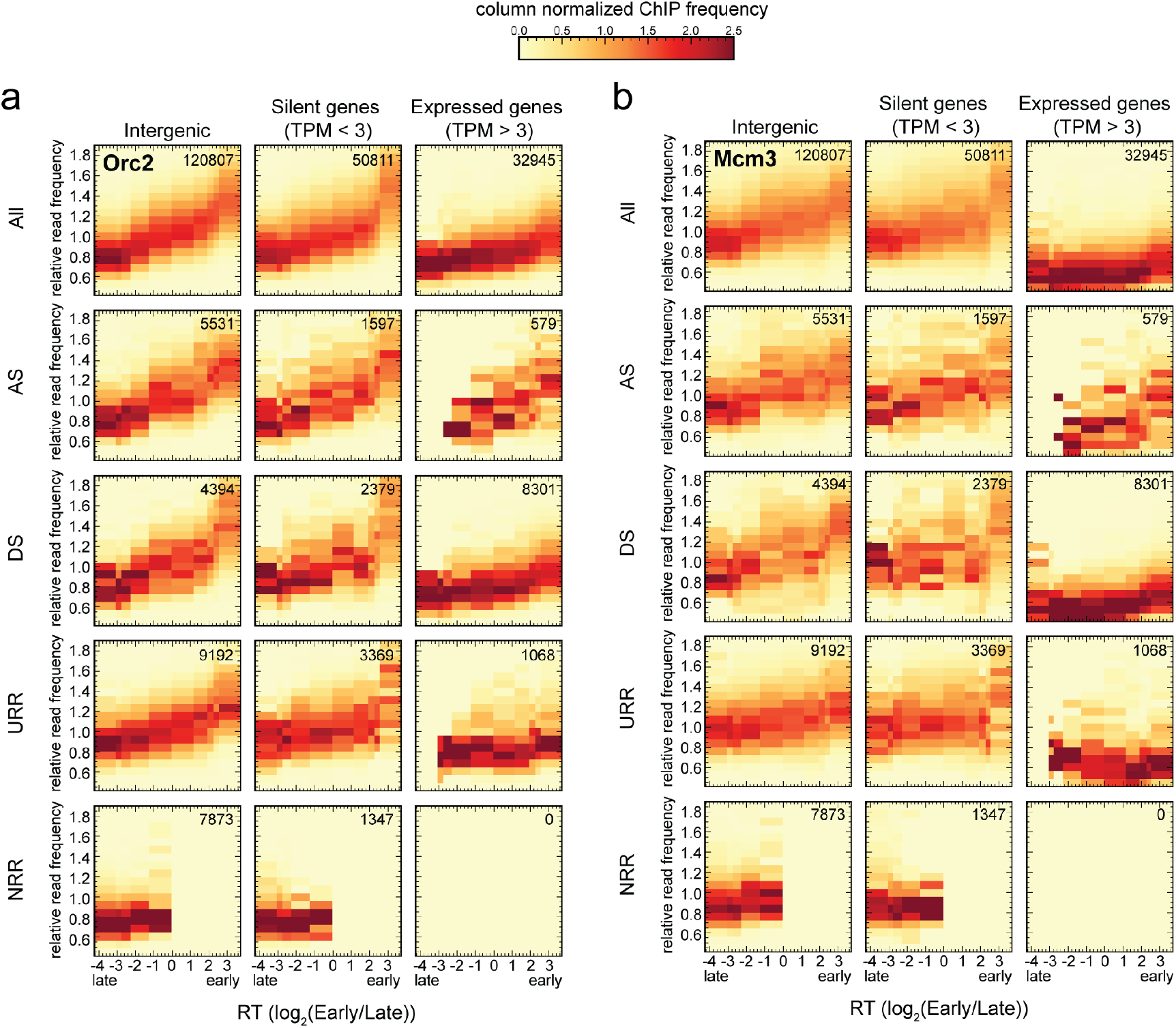
ORC/MCM levels correlate with RT and transcriptional activity but are otherwise homogeneously distributed along the genome and uncorrelated to RFD patterns (see also Suppl. Fig. 7). a-b) 3×5 panel of 2D histograms representing Orc2 (a) and Mcm3 (b) ChIP relative read frequency vs. RT (average log2(Early/Late) over 100 kb binned according to the decile of RT distribution). The analysis was performed in 10 kb windows. ChIP relative read frequencies are normalized by column and represent the probability density function of ChIP frequency at a given replication timing. The color legend is indicated on top. The columns of each panel represent only windows present in intergenic regions (left column), silent genes (TPM < 3, middle column), and expressed genes (TPM > 3, right column). TSSs and TTSs proximal regions were not considered (see Material and Methods). The rows show either all bins (top row), AS bins (predominant replication initiation, second row), DS bins (descending segment, predominant replication termination, third row), URR bins (unidirectional replication, no initiation, no termination, fourth row) and NRR bins (null RFD regions, spatially random initiation and termination, bottom row). The number of bins per panel is indicated in each panel. See Suppl. Fig. 7 for equivalent Orc3 and Mcm7 data. Refer to Suppl. Fig. 8a for statistical significances.

Silent genes (middle column) mirror the ORC/MCM pattern of intergenic regions. Expressed genes (right column) show lower ORC/MCM frequencies than intergenic regions and silent genes, as expected. Kolmogorov-Smirnov statistics (Suppl. Fig. 8a) quantitatively show that in early- and mid-replicating regions, ORC/MCM relative frequency distributions are substantially different between expressed genes and intergenic regions as well as silent genes. In contrast, intergenic distributions in ASs and DSs are not different in most RT bins. In expressed genes, the ChIP depletion is more pronounced for MCM than ORC, as described in Fig. 3a. The dependence of the residual signal on RT is much attenuated, particularly for MCM, as expected if transcription completely removes this complex from active gene bodies.

In general, the global analysis of all windows, independent from RFD, demonstrates a genome-wide, monotonous relationship between ORC/MCM binding and RT that is only attenuated in transcribed genes. Furthermore, this analysis did not reveal any convincing differences in the levels of ORC/MCM between non-transcribed ASs, DSs, URRs, and NRRs when bins of similar RT were compared. The few (579) bins corresponding to ASs in active genes are probably misleading, as they are mainly attributable to annotation errors: the annotated gene overlapped the AS but the RNA-seq signal was confined outside the AS (Petryk et al., 2016). Note that NRRs are confined to late RT segments while URRs are enriched in mid-RT segments as previously noted (Petryk et al., 2016) and confirmed in Suppl. Fig. 7c.

Strictly speaking, the slope of a RFD segment is proportional to the difference between the density of initiation and termination events within the segment (Audit et al., 2013). Therefore, we cannot exclude delocalized initiation events within DSs, which explains why they were not significantly depleted in ORC/MCM compared to ASs (Suppl. Fig. 8a). In contrast, we can mostly exclude initiation events within URRs, although their ORC/MCM densities were not significantly lower than in ASs, which are bona fide IZs, while NRRs, which presumably support spatially stochastic initiation present a smaller ORC density (Suppl. Fig. 8a).

Taken together, these results suggest that the density of ORC/MCM is not a reliable predictor of initiation probability, even though ORC density (and to a lesser extent MCM density) is well correlated with RT. Our results suggest that potential origins are widespread through the genome but that additional genetic or epigenetic factors regulate whether and when they fire.

### Late replicating non-genic ASs and NRRs are characterized by H4K20me3

The results above revealed a gradient of ORC/MCM according to RT. To confirm this observation, we extracted early and late RTDs employing a threshold of log_2_(Early/Late) > 1.6 for early RTDs and < −2.0 for late RTDs, which resulted in 302 early RTDs covering 642.8 Mb and 287 late RTDs covering 617.4 Mb of the genome. Restricting the analysis to intergenic regions, we calculated the mean ORC/MCM relative read frequencies in early compared to late RTDs. ORC was 1.4-times enriched in early RTDs compared to late RTDs (Fig. 5a, Suppl. Fig. 8b without input-correction, and Suppl. Table 2). Mcm3 and Mcm7 levels, although showing the same tendencies, were less contrasted than ORC and more influenced by input normalization (Fig. 5a, Suppl. Fig. 8b, Suppl. Table 2).

**Figure 5:**
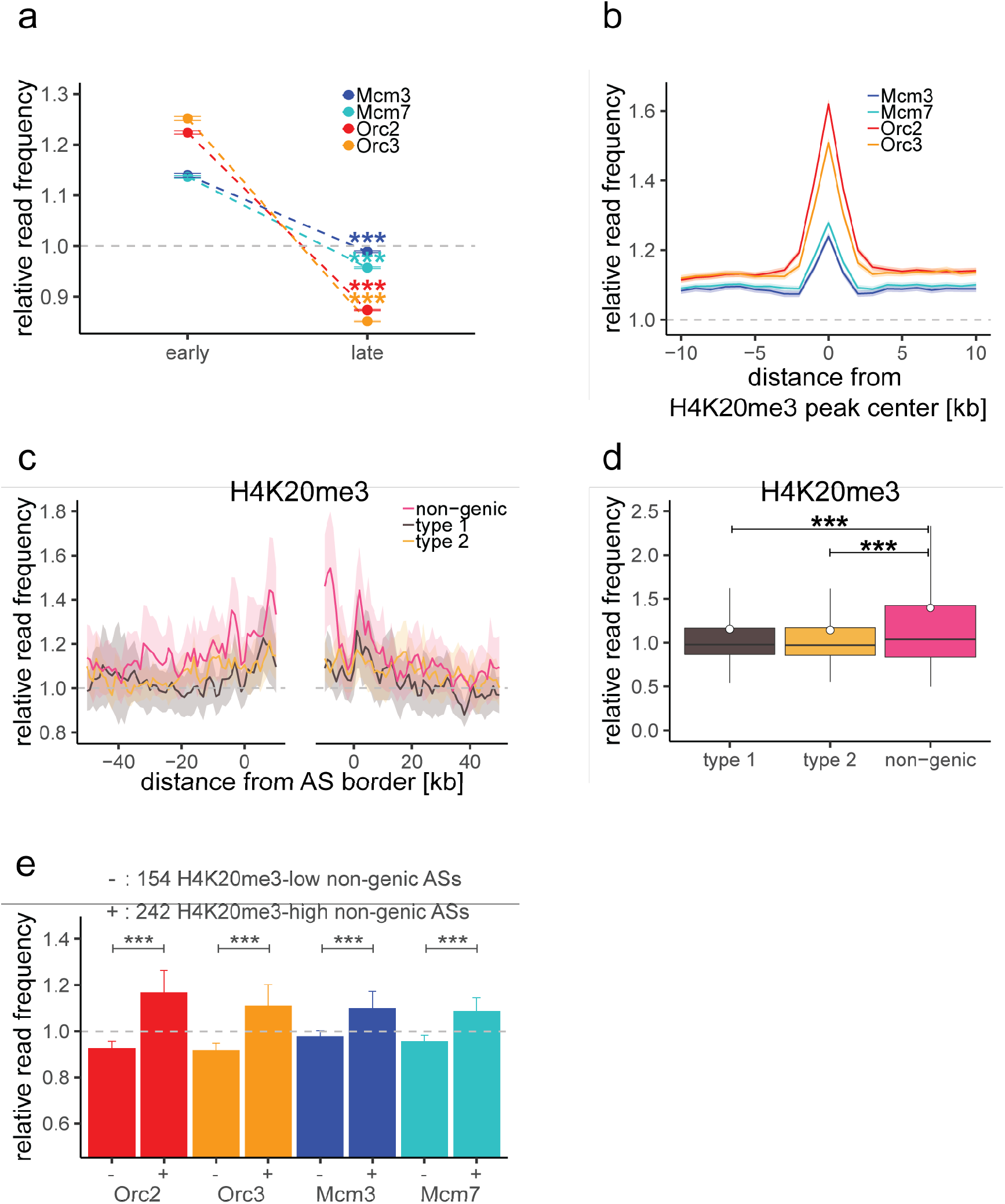
H4K20me3 selectively marks a subset of late replicating non-genic ASs. a) ORC/MCM relative read frequencies (± 2 x SEM) in early or late RTDs. Early RTDs were defined as log2(Early/Late) > 1.6; late RTDs < −2.0. The analysis was performed in 10 kb bins. Any gene ± 10 kb was removed from the analysis. Statistics were performed using one-sided t-test. b) Average ORC/MCM relative read frequencies at H4K20me3 peaks (> 1 kb). c) H4K20me3 relative read frequencies at AS borders of the different AS types. Type 2 ASs are oriented with their AS borders associated to transcribed genes at the right. Means of H4K20me3 relative read frequencies are shown ± 2 x SEM (lighter shadows). d) Boxplot representation of H4K20me3 relative read frequencies within the different AS types. Boxplot represent the mean (circle), the median (thick line), the 1st and 3rd quartile (box), the 1st and 9th decile (whiskers) of the relative read frequencies in each AS type. Statistics were performed by one-way ANOVA followed by Tukey’s post-hoc test. e) Histogram representation of mean ± 2 x SEM of ORC/MCM relative read frequencies at 242 H4K20me3-low non-genic ASs and 154 H4K20me3-high non-genic ASs. Statistics were performed using one-sided t-test. *** p < 0.001.

We recently demonstrated that H4K20me3 is involved in licensing a subset of late replicating regions, often co-localizing with H3K9me3 (Benetti et al., 2007; Brustel et al., 2017; Pannetier et al., 2008). Here, we looked further into the relation between this epigenetic mark, ORC/MCM and replication initiation. We performed ChIP for H4K20me3 and its precursor H4K20me1 in three replicates for pre-replicative G1 chromatin, validated by qPCR (Suppl. Fig. 9a (H4K20me3) and 9b (H4K20me1)). After sequencing, we performed MACS2 broad peak-detection and kept only peaks overlapping in all three samples (16852 peaks for H4K20me3 and 12264 peaks for H4K20me1, see also Suppl. Table 3 for further characterization). H4K20me3 peak sizes ranged from 200 bp to 105 kb (200 bp to 183 kb for H4K20me1, Suppl. Table 3, Suppl. Fig. 9c). When calculating ORC/MCM coverage at H4K20me3/me1 peaks > 1 kb (12251/6277 peaks, respectively), we predominantly detected ORC, but also some MCM enrichment at H4K20me3 sites (Fig. 5b). By contrast, H4K20me1 peaks were not enriched in both ORC and MCM (Suppl. Fig. 9d).

We calculated H4K20me3 coverage at the different AS types and specifically detected an increased H4K20me3 signal in non-genic ASs, representing the first histone modification characterizing late replicating ASs (Fig. 5c and 5d). Starting from 506 non-genic ASs, we extracted two subsets of 154 and 242 non-genic ASs where H4K20me3 relative read frequencies were respectively above the mean genome value by more than 1.5x standard deviation or below the genome mean value. We found that ORC/MCM were enriched at the H4K20me3-high subgroup compared to the H4K20me3-low subgroup (Fig. 5e). These results suggest that the presence of H4K20me3 at transcriptionally independent, non-genic ASs may contribute to specifying these regions as Ma-IZs at the origin-licensing step.

To further explore the links between H4K20me3 and replication initiation capability, we analyzed the density of this modification in genome segments of various RT, gene activity and RFD patterns (Fig. 6a), as performed for ORC/MCM in Fig. 4 and Suppl. Fig. 7d and 7e. Several interesting observations emerged from this analysis. First, the H4K20me3 level was weakly but systematically more abundant in early than in late replicating DNA. Second, the H4K20me3 level was slightly lower in transcribed genes than in the non-transcribed rest of the genome (Fig. 6a and Suppl. Fig 9e). Third, AS and DS bins showed comparable distributions of H4K20me3 levels at comparable RT and gene expression status (Fig. 6a and Suppl. Fig 9e). These results are, at first glance reminiscent of ORC/MCM distributions. However, ORC/MCM distributions are more pronounced as the RT gradient is stronger and the differences between expressed genes and non-transcribed intergenic regions and silent genes are more salient (Fig. 4 and Fig. 6a). NRRs showed a very different, broader distribution of H4K20me3 levels, including a higher proportion of windows with a strong H4K20me3 signal, especially compared to URRs (compare boxplots in Fig. 6a, Fig. 6b). A specific enrichment of H4K20me3 is therefore detected not only in late, non-genic AS segments but also in late-replicating gene deserts of null RFD, which presumably replicate by widespread and spatially random initiation.

**Figure 6:**
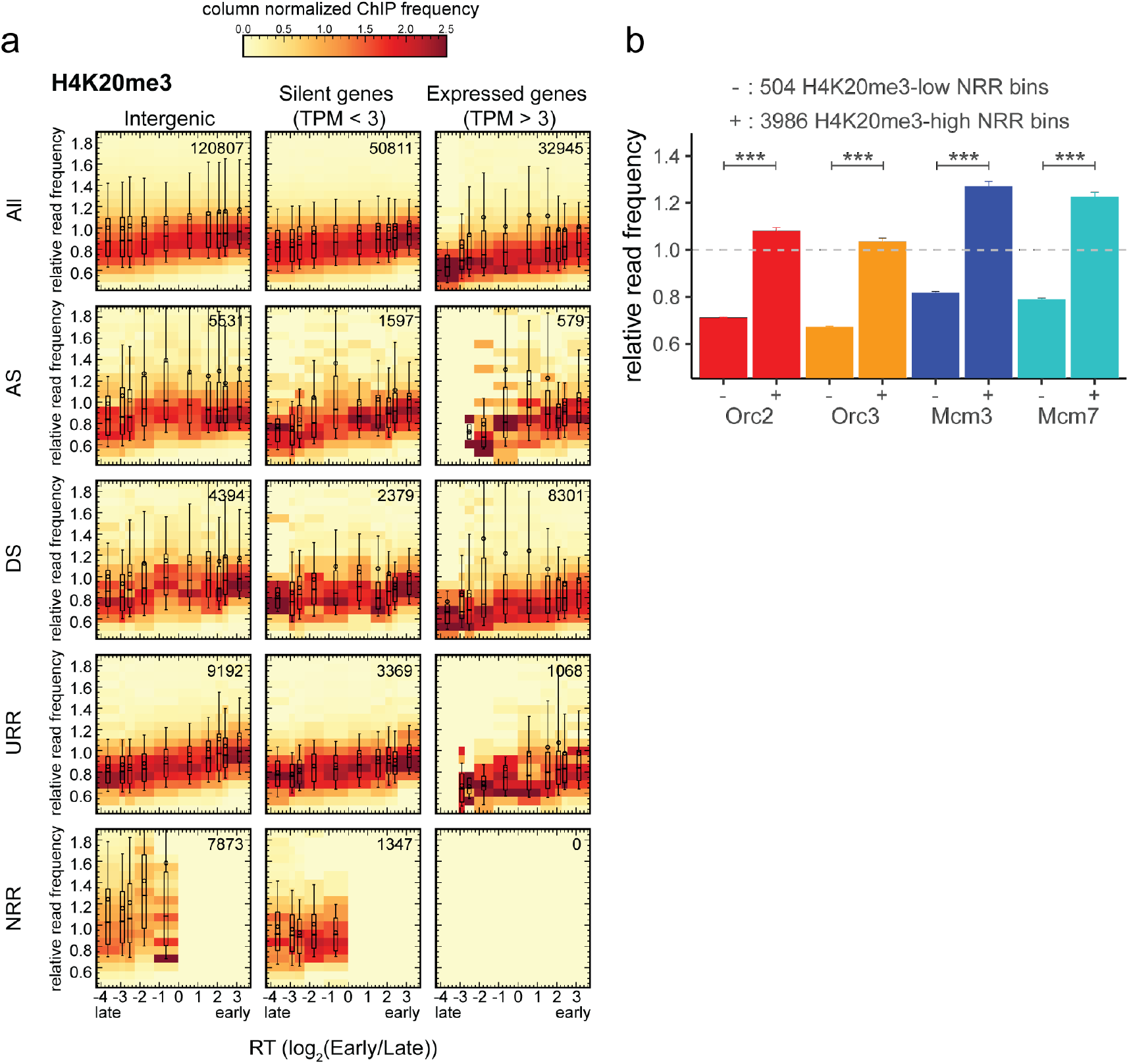
H4K20me3 is enriched in late replicating regions of null RFD (NRR). a) 3×5 panel of 2D histograms representing H4K20me3 ChIP relative read frequencies vs. RT (average log2(Early/Late) over 100 kb binned according to the decile of RT distribution). The analysis was performed in 10 kb windows. H4K20me3 relative read frequencies are normalized by column and displayed for different window categories (columns: intergenic regions, silent genes, expressed genes; rows: all bins, AS-, DS-, URR-, NRR bins) as for ORC/MCM in Fig. 4. The number of bins per panel is indicated in each panel. Superimposed boxplots represent the mean (circle), the median (thick line), the 1st and 3rd quartile (box), the 1st and 9th decile (whiskers) of the relative read frequencies in each timing bins. Refer to Suppl. Fig. 8c for statistical significances. b) Histogram representation of mean ± 2 x SEM of ORC/MCM relative read frequencies at 504 H4K20me3-low NRR 10 kb bins and 3986 H4K20me3-high NRR 10 kb bins. Statistics were performed using one-sided t-test. *** p < 0.001.

The finding that the H4K20me3 levels are more broadly distributed in NRRs suggests that ORC/MCM levels might also differ between H4K20me3-high and -low NRRs. We repeated the analysis of ORC/MCM enrichment at H4K20me3-high and - low 10 kb intergenic bins in NRRs. Similar to non-genic ASs, we also observed that ORC/MCM is more abundant at H4K20me3-high than -low bins (Fig. 5e, and 6b), supporting the hypothesis that this modification also facilitates origin licensing in these heterochromatic segments.

## Discussion

The study presented here provides a novel comprehensive genome-wide analysis of multiple pre-RC proteins, replication initiation, and replication timing in human cells. We find a widespread presence of ORC/MCM throughout the genome, with variations that only depend on RT or active transcription. ORC/MCM are depleted from transcribed genes and enriched at TSSs. ORC/MCM are more abundant in early than in late RTDs. The even distribution of ORC/MCM observed within IZs is consistent with OK-seq results suggesting a homogeneous initiation probability from any site within any given IZ. However, when RT and transcriptional effects are controlled, no significant differences in ORC/MCM densities are detected between IZs, TZs, URRs and NRRs, which sustain different patterns of origin activity. We consequently propose that potential origins, defined by loaded MCM-DHs, are widespread through the genome, but their activation in S phase is regulated by additional genetic and/or epigenetic factors. We previously reported that IZs are enriched in open chromatin marks typical of active or poised enhancers (Petryk et al., 2016), which suggests why IZs are more accessible to firing factors than flanking segments with comparable MCM-DH density. We further show that a subset of non-genic late IZs, and late, randomly replicating gene deserts, are enriched in H4K20me3, which helps to recruit ORC/MCM.

Our data suggest that transcription has both positive and negative effects on origin activity. We found that actively transcribed gene bodies flanking type 1 and type 2 ASs are depleted of ORC/MCM (Fig. 2, and Fig. 3a). We propose that active transcription removes ORC/MCM from transcribed gene bodies, which negatively affects their replication initiation capacity. MCM-DH depletion from transcribed gene bodies was previously reported in *Drosophila* (Powell et al., 2015). Experiments in yeast have shown that RNA polymerases push MCM-DHs along the DNA and redistribute them to shift replication initiation sites (Gros et al., 2015). A number of previous studies have suggested that replication does not initiate within transcribed genes (Hamlin et al., 2010; Hyrien et al., 1995; Knott et al., 2009; Macheret & Halazonetis, 2018; Martin et al., 2011; Sasaki et al., 2006). Oncogene expression, by abridging G1 phase, allows the ectopic activation of intragenic origins normally suppressed by transcription in G1 resulting in genomic instability (Macheret & Halazonetis, 2018). All these findings are consistent with an inhibitory effect of transcription on local replication initiation that is important for genome stability. Why forks emitted by ectopic, intragenic origins are more prone to genomic destabilisation than forks emitted outside, but progressing into genes, is unclear at present.

On the other hand, ORC and to a lesser degree MCM, are enriched at active TSSs. Active TSSs are regions of open chromatin structure characterized by high DNase- or MNase hypersensitivity, a hallmark of Ma-IZs (Boulos et al., 2015; Papior et al., 2012). ORC chromatin binding is known to favor open chromatin sites (Miotto et al., 2016), situating active TSSs as hotspots of ORC binding. The slope of aggregate RFD profiles around meta-TSSs is higher at TSSs than in upstream IZs, suggesting a higher probability of initiation at TSSs (Chen et al., 2019). However, a part of this effect may result from averaging multiple IZs that all end at a TSS but initiate replication at different upstream distances from the respective TSS. When individual IZs are examined, the RFD slope is not obviously increasing at the TSS (data not shown). Furthermore, the most efficient initiation sites identified by EdUseq-HU within IZs are associated with poly(dA:dT) tracts but are not enriched at TSSs (Tubbs et al., 2018). We find here that MCMs, which mark potential initiation sites, are less enriched at TSSs than ORC. Furthermore, MCMs are distributed fairly evenly upstream and downstream of transcribed gene bodies (Fig. 3a), arguing against a role for TSSs in directing MCM-DH loading specifically upstream of genes.

H4K20 methylation has multiple functions in ensuring genome integrity, such as DNA replication (Beck et al., 2012; Picard et al., 2014; Tardat et al., 2010), DNA damage repair, and chromatin compaction (Jorgensen et al., 2013; Nakamura et al., 2019; Shoaib et al., 2018), suggesting that the different functions are context-dependent and executed with different players. We previously demonstrated that H4K20me3 provides a platform to enhance licensing in late replicating heterochromatin (Brustel et al., 2017). Here, we detect both elevated ORC and MCM, when selecting for H4K20me3-enriched, non-genic ASs and NRRs (Fig 5e and 6b). Together with our previous observations (Brustel et al., 2017), we conclude that H4K20me3 is important for the licensing of at least a subset of late replicating origins in non-genic ASs and NRRs. It remains unclear whether additional chromatin features are required for licensing the remaining, H4K20me3-low non-genic ASs.

In higher eukaryotes, it is controversially discussed whether replication timing is determined by ORC (Dellino et al., 2013; Miotto et al., 2016) or MCM-DH (Das et al., 2015; Hyrien, 2016) abundance. However, potential origins are defined by assembled MCM-DHs, not by ORC. The weak correlation of MCM density with RT, and the lack of correlation with initiation capability of ASs, DSs and URRs, appear inconsistent with regulation of RT by MCM-DH density (Fig. 4b, Suppl. Fig. 7e, and Fig. 5a). We conclude that MCM-DH density therefore is not a reliable predictor of RT or RFD profiles and is unlikely to be a major determinant of RT itself. The correlation of ORC density with RT is more convincing but not necessarily causative. For example, open chromatin independently facilitates ORC binding in G1 phase and access of firing factors to MCM-DHs in S phase, resulting in correlation of ORC/MCM density with RT without implying any causal link.

The spatio-temporal replication program is relatively well conserved in consecutive replication cycles for each cell type, differs only slightly between cell lines and changes during differentiation (Hadjadj et al., 2016). Comparison with chromatin conformation capture (Hi-C) data have shown that early and late RTDs correspond to the more and less accessible compartments of the genome, respectively (Ryba et al., 2010). A higher density of chromatin interactions characterizes less accessible domains. Furthermore, both constitutive and developmentally regulated RTD boundaries align to the boundaries of topological domains, which are remarkably stable between cell types (Fragkos et al., 2015; Pope et al., 2014), and enriched in IZs (Baker et al., 2012; Petryk et al., 2016). Recently, Sima et al. used the CRISPR-Cas9 technology to identify three separate, cis-acting elements that together control the early replication time of the pluripotency-associated Dppa2/4 domain in mouse embryonic stem cells (mESCs) (Sima et al., 2019). Strikingly, these “early replication control elements” (ERCEs) are enriched in CTCF-independent Hi-C interactions and in active epigenetic marks (DNase1 HS, p300, H3K27ac, H3K4me1, H3K4me3) previously observed at OK-seq IZs (Petryk et al., 2018; Petryk et al., 2016). By mining mESC OK-seq data (Petryk et al., 2018), we found that the three ERCEs of the Dppa2/4 domain indeed fall within IZs (Suppl. Fig. 10a). Furthermore, the aggregate 1,835 ERCEs predicted genome-wide by Sima et al., from these and additional epigenetic properties of mESCs, show a significant, positive RFD shift indicative of efficient replication initiation (Suppl. Fig. 10b). This finding is confirmed in proliferating PHA-stimulated primary splenic B cells, (Suppl. Fig. 10c), attesting to the general validity of these observations.

Our data suggest that a higher ORC/MCM density *is not* a distinguishing feature of IZs from the rest of the non-transcribed genome. IZ specification therefore appears to occur at the origin activation rather than licensing step, which may be explained if the open chromatin structure found at Ma-IZs and ERCEs (Petryk et al., 2016; Sima et al., 2019) facilitates preferential accessibility to limiting firing factors during S phase (Boos & Ferreira, 2019). This is in line with the previously proposed cascade model in which replication of the human genome involves a superposition of efficient initiation at Ma-IZs identified by RFD ASs, followed by a cascade of more dispersive, less efficient origin activation along the intervening segments (Petryk et al., 2016; Wu et al., 2018).

## CONCLUSION

The mapping of ORC and MCM complexes reported here shows that in human cells, most of the genome, except transcribed genes, is licensed for replication during the G1 phase of the cell cycle. ORC/MCM are thereby more enriched in early than in late RTDs (Fig. 7a). Only a fraction of MCM-DHs is selected for initiation during S phase. Open chromatin marks define preferential IZs, often but not always circumscribed by active genes (Fig. 7b). Once forks emanate from Ma-IZs within an RTD, a cascade of replication activation may take place dispersedly between IZs due to the omnipresence of MCM-DHs (Fig. 7b). The preferential binding of ORC in G1, and preferential access of firing factors in S to open chromatin, appears sufficient to explain why ORC/MCM levels correlate with RT. Replication licensing of heterochromatic gene deserts is, for example facilitated by H4K20me3, which helps ORC and MCM-DH recruitment and supports origin activity in these less accessible chromatin segments (Fig. 7c). The identification of ERCEs supports the hypothesis that combinations of additional chromatin and DNA features regulate the probability of origin activation. This provides organizational links between active transcription and replication initiation, operating during origin licensing and activation, which facilitate the timely activation of appropriate replication origins for genome stability during programmed development as well as altered gene expression patterns caused by environmental cues.

**Figure 7:**
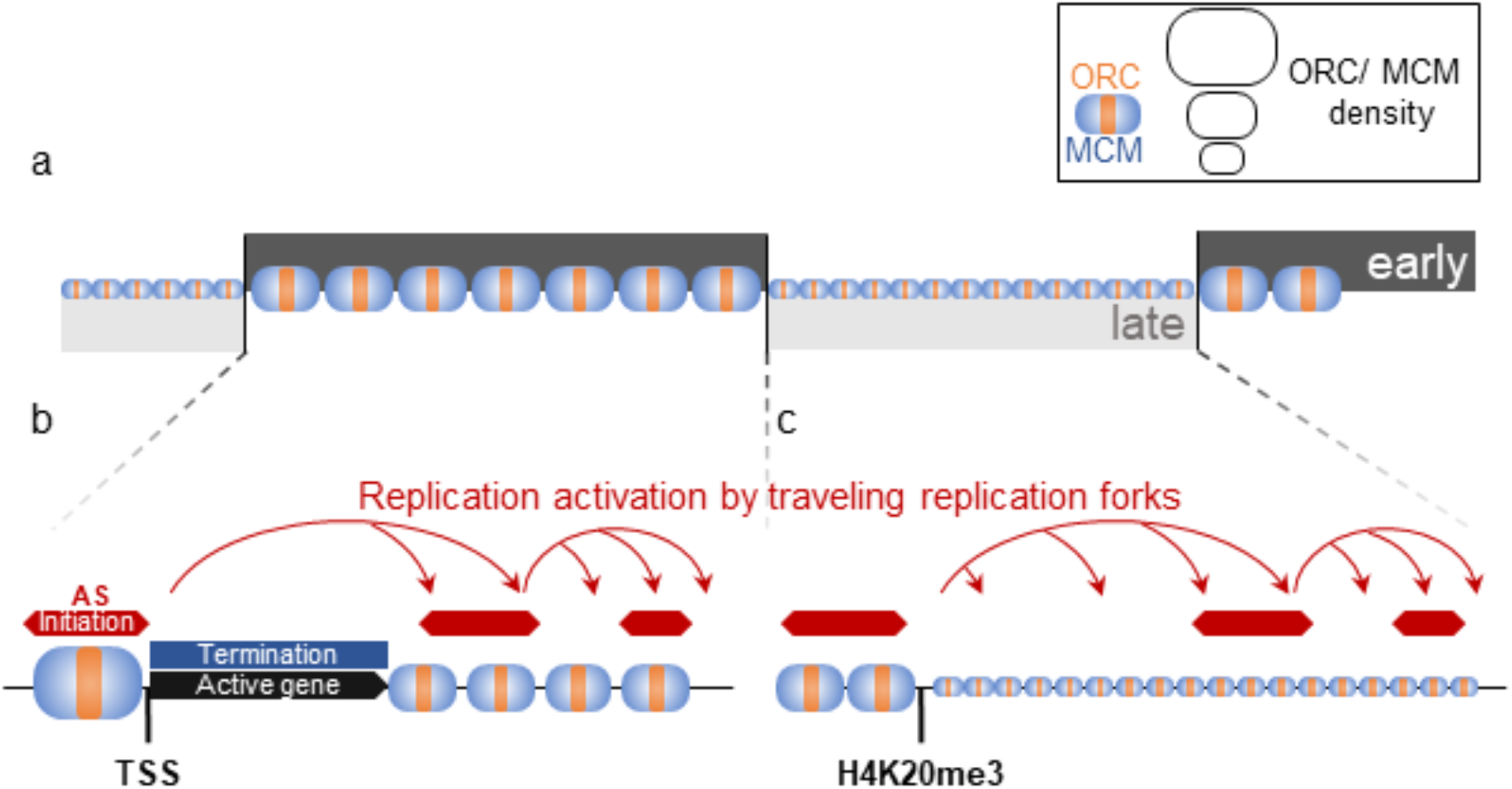
Model for replication organization in higher eukaryotes. a) Replication is organized in large segments of constant replication timing (early RTD, dark grey; late RTD, light grey), (Marchal et al., 2019). While we observe a homogeneous pattern of ORC (orange) and MCM (blue) throughout the genome, the enrichment levels of ORC/MCM were higher in early RTDs compared to late RTDs. b) Early RTDs are amongst other characterized by active transcription. ORC/MCM are locally highly enriched at active TSS. However, actively transcribed gene bodies (black) are deprived of ORC/MCM, often correlating with replication termination (blue). Besides TSSs, we find ORC/MCM stochastically distributed along intergenic regions. We hypothesize that traveling replication forks trigger activation of replication in a cascade (red arrows). c) In gene deprived and transcriptionally silent late replicating heterochromatin, we detected homogeneous ORC/MCM distribution at generally lower levels. H4K20me3 is present at late replicating non-genic ASs and NRRs and leads to enhanced ORC/MCM binding, linking this histone mark to replication activation in heterochromatin.

## Material and Methods

### Cell culture

Raji cells (ATCC) were cultivated at 37°C and 5% CO2 in RPMI 1640 (Gibco, Thermo Fisher, USA) supplemented with 8% FCS (Lot BS225160.5, Bio&SELL, Germany), 100 Units/ml Penicillin, 100 µg/ml Streptomycin (Gibco, Thermo Fisher, USA), 1x MEM non-essential amino acids (Gibco, Thermo Fisher, USA), 2 mM L-Glutamin (Gibco, Thermo Fisher, USA), and 1 mM Sodium pyruvate (Gibco, Thermo Fisher, USA).

### RNA extraction, sequencing, TPM calculation

RNA was extracted from 3 x 10^5^ Raji cells using Direct-zolTM RNA MiniPrep kit (Zymo Research) according to manufacturers’ instructions. RNA quality was confirmed by Bioanalyzer RNA integrity numbers between 9.8 and 10 followed by library preparation (Encore Complete RNA-Seq Library Systems kit (NuGEN)). Single-end 100 bp sequencing was performed by Illumina HiSeq 1500 to a sequencing depth of 25 million reads. The reads were mapped to hg19 genome using Tophat2 and assigned to annotated genes (HTSeq-count) (Anders et al., 2015; Kim et al., 2013). TPM values (Transcripts per kilobase per million reads) were calculated for each sample (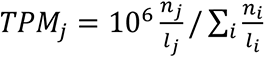 where *n*_1_ is the number of reads that map to gene *i* whose total exon length expressed in kb is *l_i_*) as previously described (Wagner et al., 2012).

### Replication fork directionality profiling using OK-seq method in Raji

Raji OK-seq was recently published as part of Wu *et al*. and is available from the European Nucleotide Archive under accession number PRJEB25180 (see data access section) (Wu et al., 2018). Reads > 10 nt were aligned to the human reference genome (hg19) using the BWA (version 0.7.4) software with default parameters (Li & Durbin, 2009). We considered uniquely mapped reads only and counted identical alignments (same site and strand) as one to remove PCR duplicate reads. Five biological replicates were sequenced providing a total number of 193.1 million filtered reads (between 19.1 and 114.1 million reads per replicate). RFD was computed as 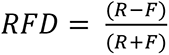, where “R” (resp. “F”) is the number of reads mapped to the reverse (resp. forward) strand of the considered regions. RFD profiles from biological replicates were highly correlated, with Pearson correlation computed in 50 kb non-overlapping windows with > 100 mapped reads (R+F) ranging from 0.962 to 0.993. Reads from the 5 replicate experiments were pooled together for further analyses.

### Determining regions of ascending, descending and constant RFD

RFD profiling of 2 human cell lines revealed that replication primarily initiates stochastically within broad (up to 150 kb) zones and terminates dispersedly between them (Petryk et al., 2016). These initiation zones correspond to quasi-linear ascending segments (ASs) of varying size and slope within the RFD profiles. As previously described for mean replication timing profiles analysis (Audit et al., 2013; Baker et al., 2012), we determined the smoothed RFD profile convexity from the convolution with the second derivative of the Gaussian function of standard deviation 32 kb. 4891 ASs were delineated as the regions between positive and negative convexity extrema of large amplitude. The amplitude threshold was set in a conservative manner in order to mainly detect the most prominent IZs described by Petryk et al. (2016) and avoid false positives. Descending segments (DSs) were detected symmetrically to ASs, as regions between negative and positive convexity extrema using the same threshold. Noting *pos_5’* and *pos_3’* the location of the start and end position of an AS or DS segment, each segment was associated to its size *pos_3’-pos_5’* and the RFD shift across its length: ΔRFD = RFD (*pos_5’*) – RFD (*pos_3’*). DS segments were less numerous (2477 vs 4891) and on average larger (126 kb vs 38.8 kb) than AS segments as expected and presented a smaller average RFD shift (|ΔRFD|=0.69 vs 0.83).

Initial RFD profiling in human also revealed regions of unidirectional fork progression and regions of null RFD where replication is bidirectional. Unidirectional replicating regions (URRs) were delineated as regions where |ΔRFD|>0.8 homogeneously over at least 300 kb (401 regions of mean length 442 kb covering 177 Mb). Null RFD regions (NRRs) were delineated as regions where |ΔRFD|<0.15 homogeneously over at least 500 kb (127 regions of mean length 862 kb covering 110 Mb). Thresholds were set in a conservative manner to avoid false positive, in particular not to confuse RFD zero-crossing segments with NRR.

### Centrifugal elutriation and flow cytometry

For centrifugal elutriation, 5 x 10^9^ exponentially growing Raji cells were harvested, washed with PBS and resuspended in 50 ml RPMI 1680, 8% FCS, 1mM EDTA, 0.25 U/ml DNaseI (Roche, Germany). Concentrated cell suspension was passed through 40 µm cell strainer and injected in a Beckman JE-5.0 rotor with a large separation chamber turning at 1500 rpm and a flow rate of 30 ml/min controlled by a Cole-Parmer Masterflex pump. While rotor speed was kept constant, 400 ml fractions were collected at increasing flow rates (40, 45, 50, 60, 80 ml/min). Individual fractions were quantified, 5 x 10^6^ cells washed in PBS, ethanol fixed, RNase treated and stained with 0.5 mg/ml Propidium Iodide. DNA content was measured using the FL2 channel of FACSCalibur^TM^ (BD Biosciences, Germany). Remaining cells were subjected to chromatin cross-linking.

### Chromatin cross-linking with formaldehyde

Raji cells were washed twice with PBS, resuspended in PBS to a concentration of 2 x 10^7^ cells/ml and passed through 100 µm cell strainer (Corning Inc., USA). Fixation for 5 min at room temperature was performed by adding an equal volume of PBS 2% methanol-free formaldehyde (Thermo Scientific, USA, final concentration: 1% formaldehyde) and stopped by the addition of glycine (125 mM final concentration). After washing once with PBS and once with PBS 0.5% NP-40, cells were resuspended in PBS containing 10% glycerol, pelleted and snap frozen in liquid nitrogen.

### Cyclin Western Blot

Cross-linked samples were thawed on ice, resuspended in LB3+ sonication buffer containing protease inhibitor and 10 mM MG132. After sonicating 3 x 5 min (30 sec on, 30 sec off) using Bioruptor in presence of 212-300 µm glass beads, samples were treated with 50 U Benzonase for 15 min at room temperature and centrifuged 15 min at maximum speed. 50 µg protein lysates were loaded on 10% SDS-polyacrylamid gel (Cyclin A1/A2, Cyclin B1), or 12.5%-15% gradient gel (H3S10P). Cyclin A1/A2 (Abcam, ab185619), Cyclin B1 (Abcam, ab72), H3S10P (Cell signaling, D2C8) antibodies were used in 1:1000 dilutions, GAPDH (clone GAPDH3 10F4, rat IgG2c; Monoclonal Antibody Core Facility, Helmholtz Center München) was diluted 1:50. HRP-coupled secondary antibodies were used in 1:10000 dilutions. Detection was done using ECL on CEA Blue Sensitive X-ray films.

### Chromatin sonication

Cross-linked cell pellets were thawed on ice, resuspended LB3+ buffer (25 mM HEPES (pH 7.5), 140 mM NaCl, 1 mM EDTA, 0.5 mM EGTA, 0.5% Sarkosyl, 0.1% DOC, 0.5% Triton-X-100, 1X protease inhibitor complete (Roche, Germany)) to a final concentration of 2 x 10^7^ cells/ml. Sonication was performed in AFA Fiber & Cap tubes (12×12 mm, Covaris, Great Britain) at an average temperature of 5°C at 100W, 150 cycles/burst, 10% duty cycle, 20 min using the Covaris S220 (Covaris Inc., UK) resulting in DNA fragments of 100-300bp on average.

### Chromatin immunoprecipitation and qPCR quality control

Sheared chromatin was pre-cleared with 50 µl protein A Sepharose 4 Fast Flow beads (GE Healthcare, Germany) per 500 µg chromatin for 2h. 500 µg chromatin (or 250 µg for histone methylation) were incubated with rabbit anti-Orc2, anti-Orc3, anti-Mcm3, anti-Mcm7 (Papior et al., 2012), mouse anti-H4K20me1 (Diagenode, MAb-147-100), rabbit anti-H4K20me3 (Diagenode, MAb-057-050), or IgG isotype controls for 16h at 4°C. BSA-blocked protein A beads (0.5 mg/ml BSA, 30 µg/ml salmon sperm, 1X protease inhibitor complete, 0.1% Triton-X-100 in LB3(-) buffer (without detergents)) were added (50 µl/500 µg chromatin) and incubated for at least 4h on an orbital shaker at 4°C. Sequential washing steps with RIPA (0.1% SDS, 0.5% DOC, 1% NP-40, 50 mM Tris (pH 8.0), 1 mM EDTA) 150mM NaCl, RIPA-300 mM NaCl, RIPA-250 mM LiCl buffer, and twice in TE (pH 8.0) buffer were performed. Immunoprecipitated chromatin fragments were eluted from the beads by shaking twice at 1200 rpm for 10 min at 65°C in 100µl TE 1% SDS. The elution was treated with 80 µg RNAse A for 2h at 37°C and with 8 µg proteinase K at 65°C for 16h. DNA was purified using the NucleoSpin Extract II Kit. Quantitative PCR analysis of the EBV *oriP* DS element (for pre-RC ChIP), or H4K20me1 and - me3 positive loci were performed using the SYBR Green I Master Mix (Roche) and the Roche LightCycler 480 System. Oligo sequences for qPCR were DS_fw: AGTTCACTGCCCGCTCCT, DS_rv: CAGGATTCCACGAGGGTAGT, H4K20me1positive_fw: ATGCCTTCTTGCCTCTTGTC, H4K20me1positive_rv: AGTTAAAAGCAGCCCTGGTG, H4K20me3positive_fw: TCTGAGCAGGGTTGCAAGTAC, H4K20me3positive_rv: AAGGAAATGATGCCCAGCTG. Chromatin fragment sizes were verified by loading 1-2 µg chromatin on a 1.5% agarose gel. Samples were quantified using Qubit HS dsDNA.

### ChIP-sample sequencing

ChIP sample library preparations from > 4 ng of ChIP-DNA was performed using Accel-NGS® 1S Plus DNA Library Kit for Illumina (Swift Biosciences). 50 bp single-end sequencing was done with the Illumina HiSEQ 1500 sequencer to a sequencing depth of ∼ 70 million reads. Fastq-files were mapped against the human genome (hg19, GRCh37, version 2009), extended for the EBV genome (NC007605) using bowtie (v1.1.1), (Langmead et al., 2009). Sequencing profiles were generated using deepTools’ bamCoverage funtion using reads extension to 200 bp and reads per genomic content (RPGC) normalization (Ramirez et al., 2014). Visualization was performed in UCSC Genome Browser (http://genome.ucsc.edu) and the Integrated Genome Browser (Kent et al., 2002).

For H4K20me1 and -me3 ChIP-seq data, MACS2 peak-calling (Zhang et al., 2008) was performed using the broad setting and overlapping peaks in three replicates were retained for further analyses.

### Binning approach and normalization

All data processing and analysis steps were performed in R (v.3.2.3) and numpy (v.1.18.5) python library, visualizations were done using the ggplot2 (v3.1.0) package (R_Core_Team, 2018) and matplotlib (v.3.2.3) python library. The numbers of reads were calculated in non-overlapping 1 or 10 kb bins and saved in bed files for further analysis. To combine replicates, their sum per bin was calculated (= read frequency). To adjust for sequencing depth, the mean frequency per bin was calculated for the whole sequenced genome and all bins’ counts were divided by this mean value resulting in the normalized read frequency. To account for variations in the input sample, we additionally divided by the normalized read frequency of the input, resulting in the relative read frequency. When aggregating different loci, input normalization was performed after averaging. This resulted in relative read frequency ranging from 0 to ∼30. Pair-wise Pearson correlations of ORC/MCM samples were clustered by hierarchical clustering using complete linkage clustering.

### Relation of ChIP relative read frequencies to DNase hypersensitivity

The ENCODE ‘DNase clusters’ track wgEncodeRegDnaseClusteredV3.bed.gz (03.12.2017) containing DNase hypersensitive sites from 125 cell lines were retrieved from (Thurman et al., 2012). Bins overlapping or not with HS sites larger than 1 kb were defined and the respective ChIP read frequency assigned for comparison.

### Comparison of ChIP relative read frequencies to replication data

ASs were aligned on their left (5’) and right (3’) borders. Mean and standard error of the mean (SEM) of relative read frequencies of aligned 1 kb bins were then computed to assess the average ChIP signal around the considered AS borders 50 kb away from the AS to 10 kb within the AS. To make sure bins within the ASs were closer to the considered AS border than to the opposite border, only ASs of size >20 kb were used (3247/4891). We also limited this analysis to ASs corresponding to efficient initiation zones by requiring ΔRFD > 0.5, filtering out a further 290 lowly efficient ASs.

In order to interrogate the relationship between ASs and transcription, we compared the results obtained for different AS groups: 506 ASs were classified as non-genic AS when the AS locus extended 20-kb at both ends did not overlap any annotated gene; the remaining 2451 ASs were classified as genic ASs. From the latter group, 673 ASs were classified as type 1 ASs when both AS borders where flanked by at least one actively transcribed genes (distance of both AS borders to the closest transcribed (TPM > 3) gene body was < 20 kb), and 1026 ASs were classified as type 2 ASs when only one AS border was associated to a transcribed gene (see also Table 1).

In order to assess the role of H4H20me3 mark on AS specification, we also classified non-genic ASs depending on their input normed H4K20me3 relative read frequency. We grouped the non-genic ASs where the H4K20me3 relative read frequency was above the genome mean value by more than 1.5 standard deviation (estimated over the whole genome) and the non-genic ASs where the H4K20me3 relative read frequency was below the genome mean value. This resulted in 154 non-genic ASs with H4K20me3 signal significantly higher than genome average and 242 non-genic ASs with H4K20me3 signal lower than genome average.

A similar selection was performed on fully intergenic 10 kb windows within NRRs (as done above using the mean and standard deviation of H4K20me3 relative read frequency estimated on all fully intergenic 10 kb windows). This resulted in 504 and 3986 windows with high and low H4K20me3 signal, respectively.

### Comparison of ChIP relative read frequencies to transcription data

Gene containing bins were determined and overlapping genes removed from the analysis. For cumulative analysis, we only worked with genes larger 30 kb, and assigned the gene expression levels in TPM accordingly. Genes were either aligned at their transcriptional start site (TSS) or their transcriptional termination site (TTS) and the corresponding ChIP read frequencies were calculated in a 30 kb window around the site.

### Comparison of ChIP relative read frequencies to replication timing

For identification of RTDs in Raji cells, we used the early to late replication timing ratio determined by Repli-seq (Sima et al., 2018). We directly worked from the precomputed early to late log-ratio from supplementary file GSE102522_Raji_log2_hg19.txt downloaded from GEO (accession number GSE102522). The timing of every non-overlapping 10 kb bin was calculated as the averaged log_2_(Early/Late) ratio within the surrounding 100 kb window. Early RTDs were defined as regions where the average log-ratio > 1.6 and late RTDs as regions where the average log-ratio < -2.0. These thresholds resulted in 1648 early RTDs, ranging from 10 to 8940 kb in size, with a mean size of 591 kb, while we detected 2046 late RTDs in sizes from 10 to 8860 kb, averaging at 470 kb. These RTDs were used to classify ChIP read relative frequencies calculated in 10 kb bins as early or late replication timing. Bins overlapping gene extended by 10 kb on both sides were removed from the analysis to avoid effects of gene activity on ChIP signals.

### Comparison of ChIP relative read frequencies distributions at different replication timing depending on transcriptional and replicative status

All non-overlapping 10 kb windows were classified as intergenic if closest genes were more than 5 kb away, as belonging to a silent (resp. expressed) gene body if the window was inside a gene with TPM<3 (resp. TPM>3) and at more than 3 kb of gene borders, otherwise windows were disregarded. This made sure that specific ChIP signal at gene TSS and TTS were not considered in the analysis. Using the 3 window categories, we computed the 2D histograms of ChIP relative read frequencies vs replication timing in intergenic, silent and expressed gene bodies. We used 10 timing bins corresponding to the deciles of the whole genome timing distribution. For each timing bin, the histogram counts were normalized so as to obtain an estimate of the probability distribution function of the ChIP signal at the considered replication timing. The analysis was reproduced after restricting for windows fully in (i) AS segments (size > 20 kb, ΔRFD > 0.5), (ii) DS segments (size > 20 kb, ΔRFD < -0.5), (iii) URRs and (iv) NRRs.

### Statistics

Statistical analyses were performed in R using one-sided t-test with Welch correction and 95% confidence interval or one-way ANOVA followed by Tukey’s multiple comparisons of means with 95% family-wise confidence level, if appropriate. Comparison between ChIP signal distribution observed in two situations were performed computing the 2 sample Kolmogorov–Smirnov statistics *D_KS_* using SciPy (v.1.5.0) statistical library and correcting for sample sizes by reporting 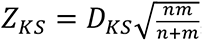, where *n* and *m* are the sizes of the two samples respectively.

### ERCE RFD profiles

The position of the three genetically identified ERCEs in the mESC Dppa2/4 locus and of the 1,835 predicted mESC ERCEs were downloaded from Sima et al. (Sima et al., 2019. The mESC OK-seq data were downloaded from Petryk et al. (Petryk et al., 2018), (SRR7535256), and mapped to mm10 genome Petryk et al. (Petryk et al., 2018). OK-seq data from cycling mouse B cells were downloaded from Tubbs et al. (Tubbs et al., 2018) (GSE116319). The RFD profile was computed as in Hennion et al., 2020 with 10kb binning steps. Predicted ERCE shuffling was performed using a homemade function keeping the number of ERCE constant for each chromosome and avoiding unmapped genome sequences (genome regions with more than 20 consecutives N). Aggregated average RFD profiles were centered on the ERCE and the profile’s envelopes represent the CI95 based on the mean and standard deviation at each position.

### Data access

Data have been deposited to the European Nucleotide Archive (ENA, https://www.ebi.ac.uk/ena). OK-seq data in Raji cells are available under the accession numbers PRJEB25180 (study accession) and SAMEA104651899 (sample accession, 5 replicates). Raji RNA-seq data are available under the accession number PRJEB31867 (study) and SAMEA5537240, SAMEA5537246, and SAMEA5537252 (sample accession per replicate). Raji ChIP-seq data were deposited under the accession number PRJEB32855.

## Acknowledgements

We thank Tobias Straub for initial help with bioinformatical analyses, Torsten Krude and Till Bartke for critical comments on the manuscript.

A.S. was supported by the Deutsche Forschungsgemeinschaft (SFB 1064 TP05), SPP1230 and by the HELENA graduate school of the Helmholtz Zentrum München. B.A. and O.H were supported by the Agence Nationale de la Recherche (ANR-15-CE12-0011, ANR-18-CE45-0002, ANR-19-CE12-0028) and the Fondation pour la Recherche Médicale (FRM DEI201512344404), and the Cancéropôle Ile-de-France and the INCa (PL-BIO16-302). O.H. was supported by the Ligue Nationale Contre le Cancer (Comité de Paris), the Association pour la Recherche sur le Cancer, and the program “Investissements d’Avenir” launched by the French Government and implemented by the ANR (ANR-10-IDEX-0001-02 PSL*Research University). W.H. was supported by the Deutsche Forschungsgemeinschaft (SFB1064/TP A13, SFB-TR36/TP A04), Deutsche Krebshilfe (grant number 70112875), and National Cancer Institute (grant number CA70723).

## Author contributions

N.K. and A.S. designed and N.K. performed the majority of experiments; A.B. performed the RNA-seq experiment and TPM analysis; X.W. performed OK-seq experiments, S.K. and H.B. generated the sequencing library and sequencing, W.H. designed RNA-seq experiments; O.H. developed OK-seq, B.A supervised bioinformatic analyses; N.K. L.L. and B.A. performed bioinformatic analyses; A.S. and OH proposed and designed the project and experimental systems; N.K., O.H. and A.S. wrote the manuscript with comments from L.L. and B.A.; All the authors read and approved the manuscript.

## SUPPLEMENTARY MATERIAL

**Supplementary Figure 1:**
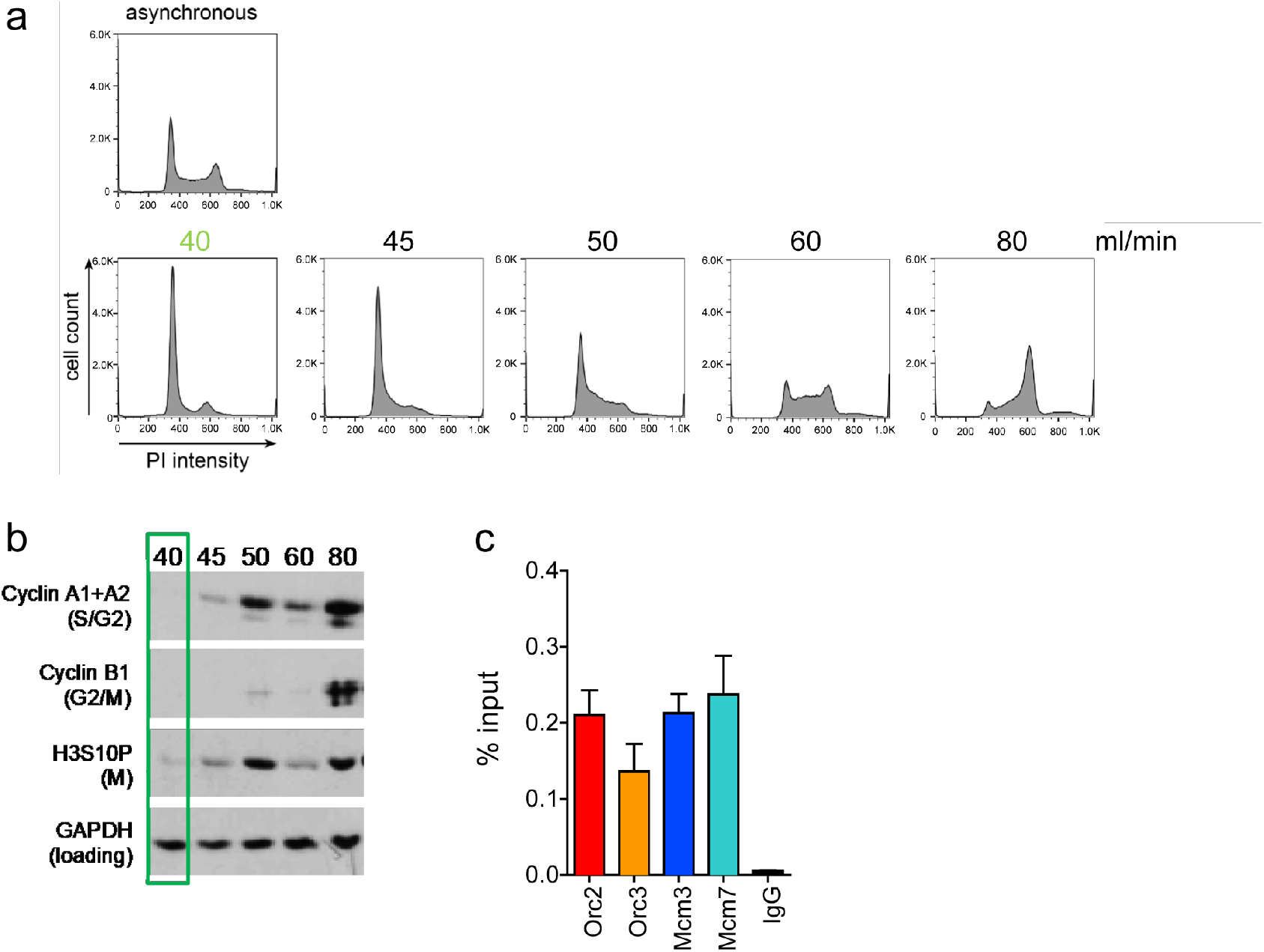
Experimental validation of cell cycle fractionation and ORC/MCM ChIP quality. a) Example DNA content (Propidium Iodide) staining followed by FACS of logarithmically growing Raji (top) cells after cell cycle fractionation by centrifugal elutriation (increasing counter flow rates indicated above each profile). b) Western Blot analyses of the single fractions detecting Cyclin A (S/G2), Cyclin B (G2/M), H3S10P (M) and GAPDH. c) qPCR validation of Orc2, Orc3, Mcm3 and Mcm7 enrichment at the EBV latent origin oriP dyad symmetry element. Representation in % input. Isotype IgG was used as control.

**Supplementary Figure 2:**
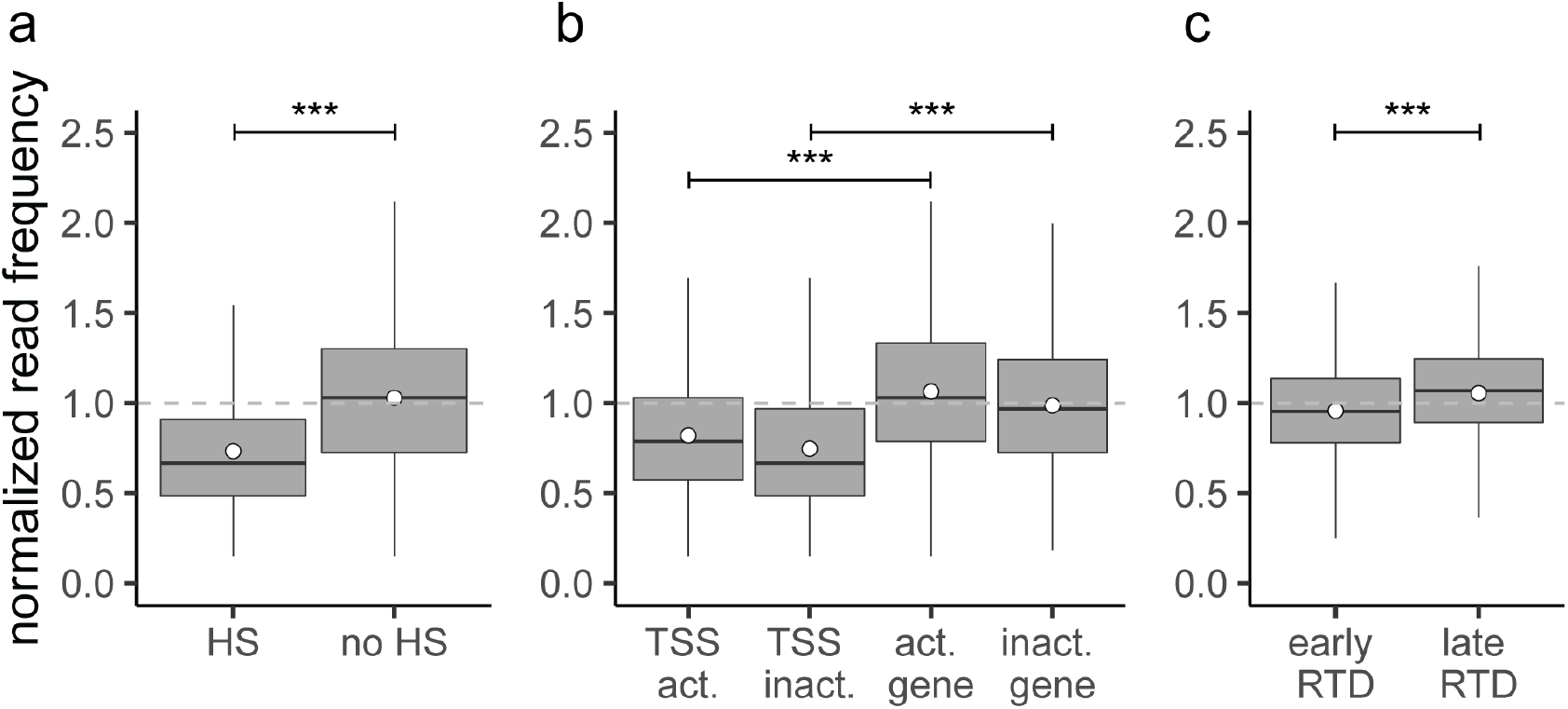
The input sequencing control is differentially represented in regions of biological function. Boxplot of normalized input read frequencies in relation to a) DNase hypersensitivity. DNase HS clusters were obtained from 125 cell lines in ENCODE, only HS sites larger 1 kb were considered. b) transcription: TSSs and gene body of active (TPM > 3) and inactive (TPM < 3) genes, and c) early or late replication timing domains (RTDs). The dashed grey horizontal line indicates read frequency 1.0 for orientation. Boxplot represent the mean (circle), the median (thick line), the 1st and 3rd quartile (box), the 1st and 9th decile (whiskers) of the relative read frequencies, without representing outliers. Statistics were performed using one-sided t-test. * p < 0.05, ** p < 0.01, *** p < 0.001.

**Supplementary Figure 3:**
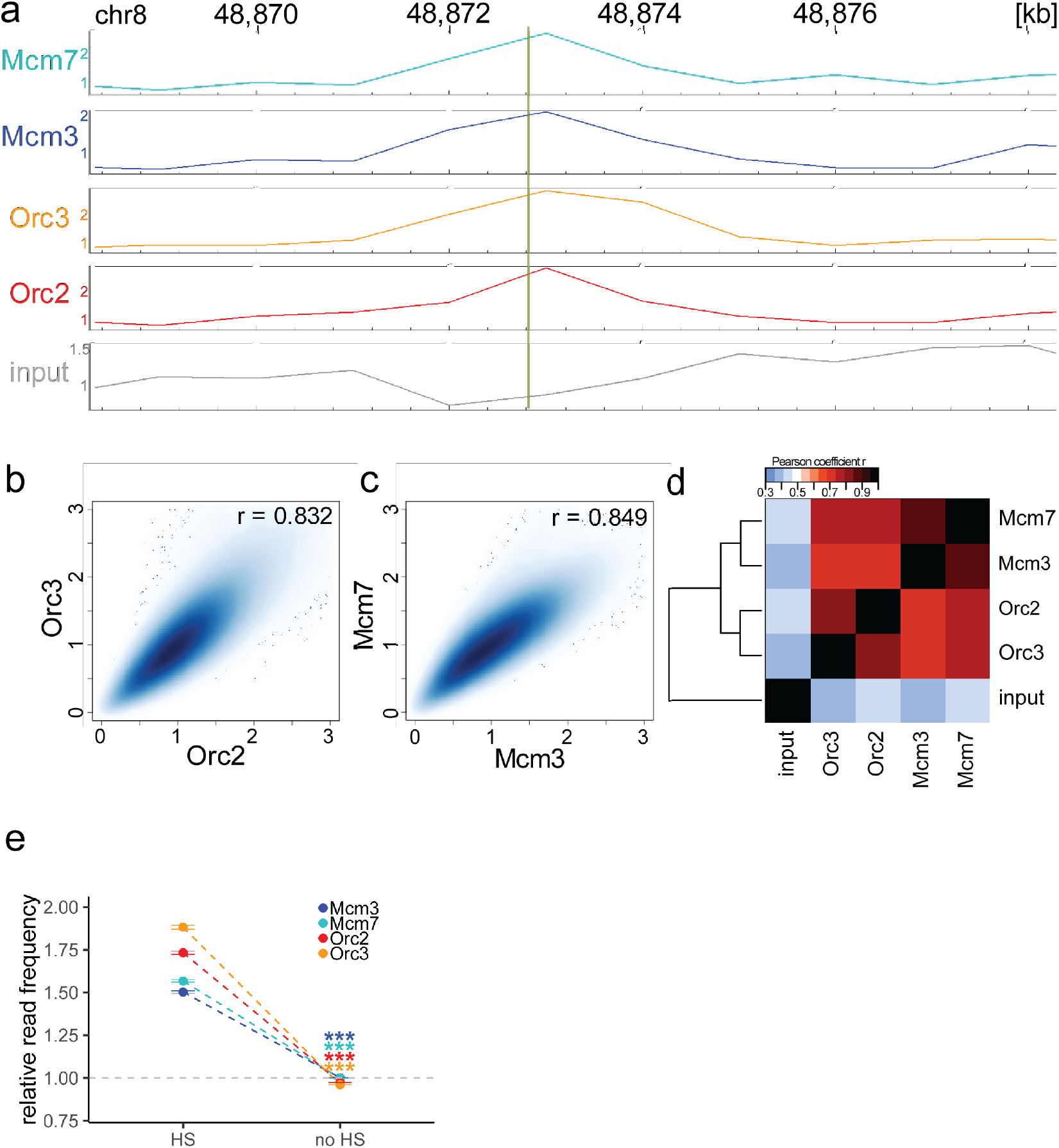
ORC/MCM enrichment at the MCM4/PRKDC origin persists without input normalization. a) The profile of ORC/MCM ChIP-seq after 1 kb binning in the same 10 kb window as Figure 1b (chr8: 48,868,314 - 48,878,313). The reads of replicates were summed and normalized by the total genome-wide ChIP read frequency. Y-axis represents the resulting normalized read frequency. b) Correlation plot between Orc2 and Orc3 normalized read frequencies in 1 kb bins. c) Correlation plot between Mcm3 and Mcm7 normalized read frequencies in 1 kb bins. d) Heatmap of Pearson correlation coefficients r between all ChIP normalized read frequencies including input in 1 kb bins. Column and line order were determined by complete linkage hierarchical clustering using the correlation distance (d = 1-r). e) ORC/MCM binding is confirmed at DNase HS sites: mean input-normalized ORC/MCM relative read frequencies (± 2 x SEM) in relation to DNase hypersensitivity. Only HS sites larger 1 kb were considered. Statistics were performed using one-sided t-test. *** p < 0.001.

**Supplementary Figure 4:**
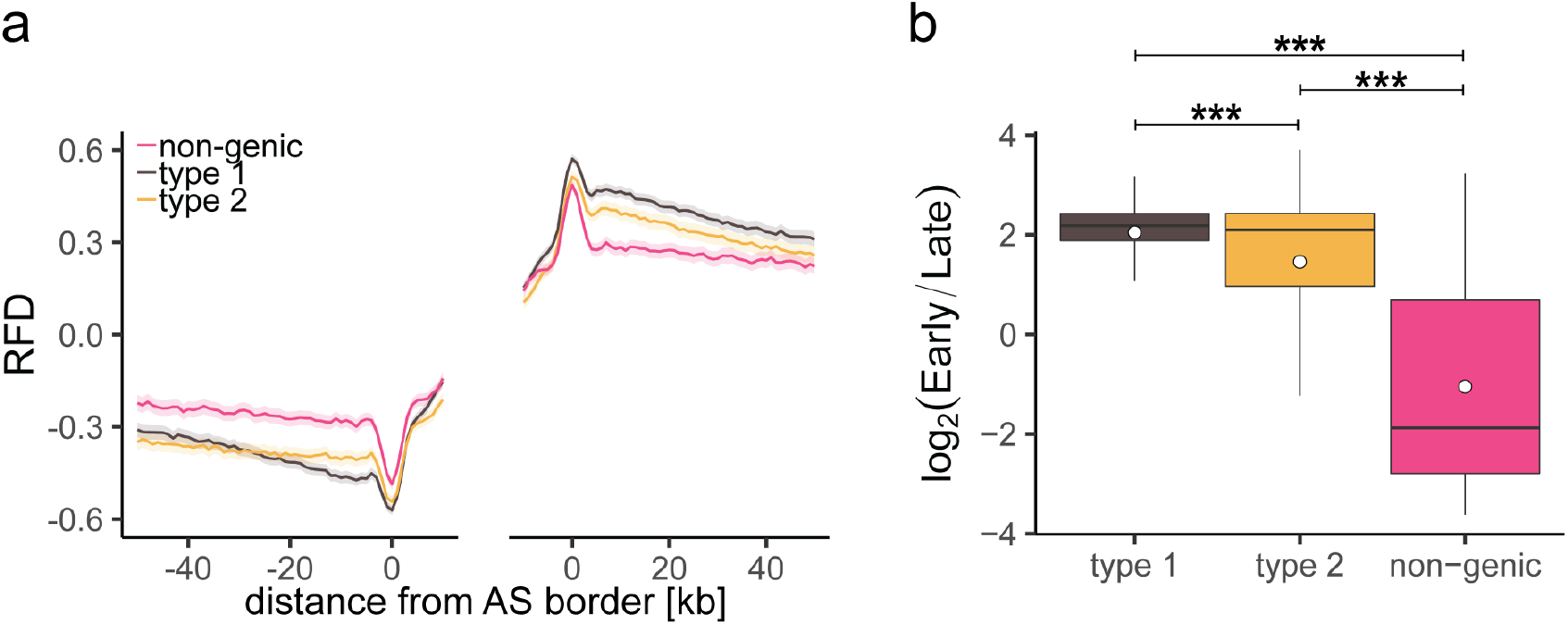
Characterization of different AS types. a) Average RFD of different AS types plotted at AS borders ± 2 x SEM (lighter shadows). b) Replication timing ratio log2(Early/Late) was assigned to type 1, type 2, and non-genic AS and represented as boxplot. Boxplot represent the mean (circle), the median (thick line), the 1st and 3rd quartile (box), the 1st and 9th decile (whiskers) of the relative read frequencies, without representing outliers. Statistics were performed by one-way ANOVA followed by Tukey’s post-hoc test. *** p < 0.001.

**Supplementary Figure 5:**
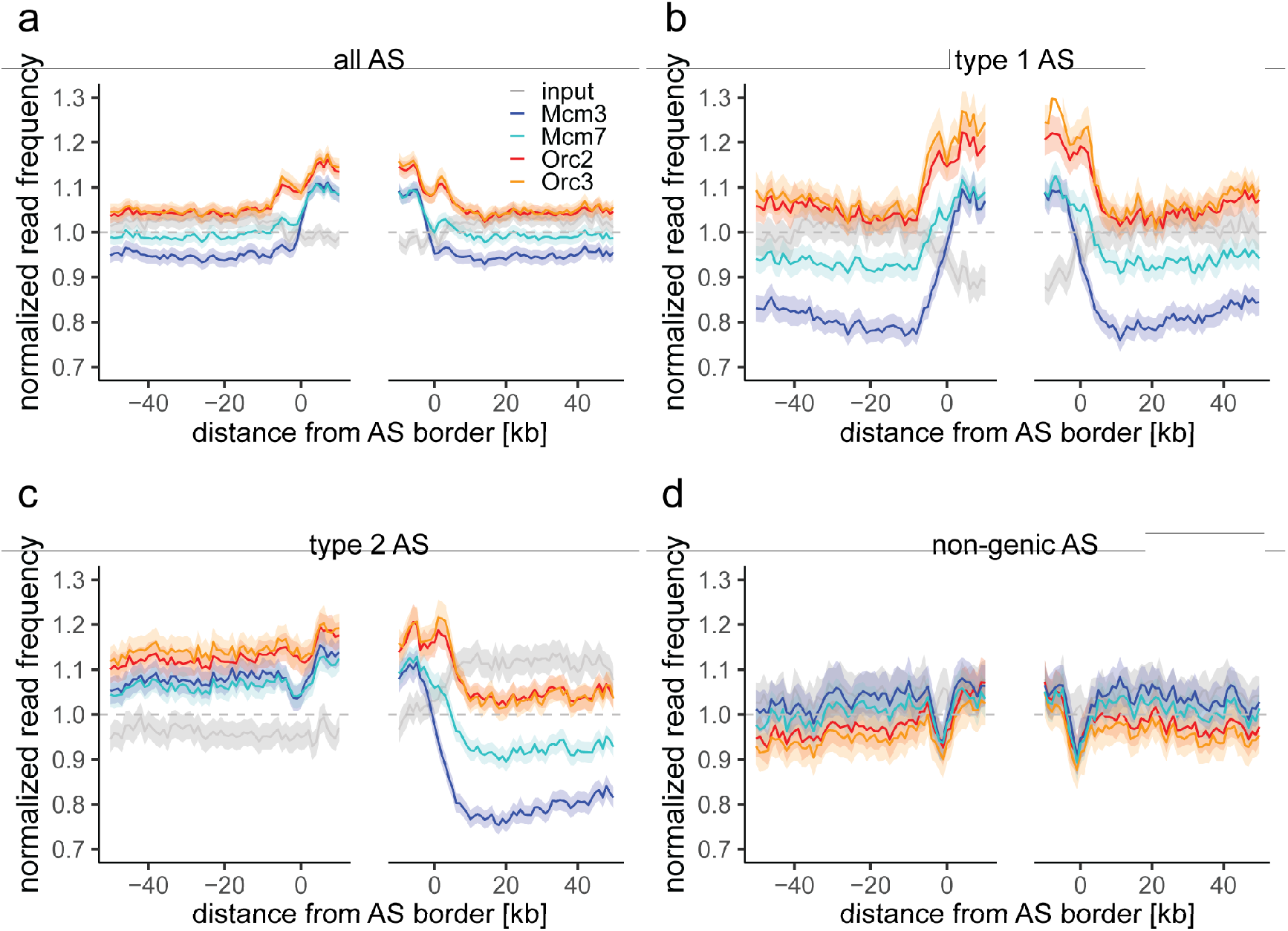
ORC/MCM enrichment within AS without input normalization. a-d) Average ChIP normalized read frequencies of Orc2, Orc3, Mcm3, Mcm7, and input at AS borders of b) all ASs (n = 2957), c) type 1 ASs with transcribed genes at both ASs borders (n = 673), d) type 2 ASs with transcribed genes oriented at their right ASs border (n = 1026), and e) non-genic ASs in gene deprived regions (n = 506). The mean of normalized read frequencies is shown ± 2 x SEM (lighter shadows). The dashed grey horizontal line indicates read frequency 1.0 for reference.

**Supplementary Figure 6:**
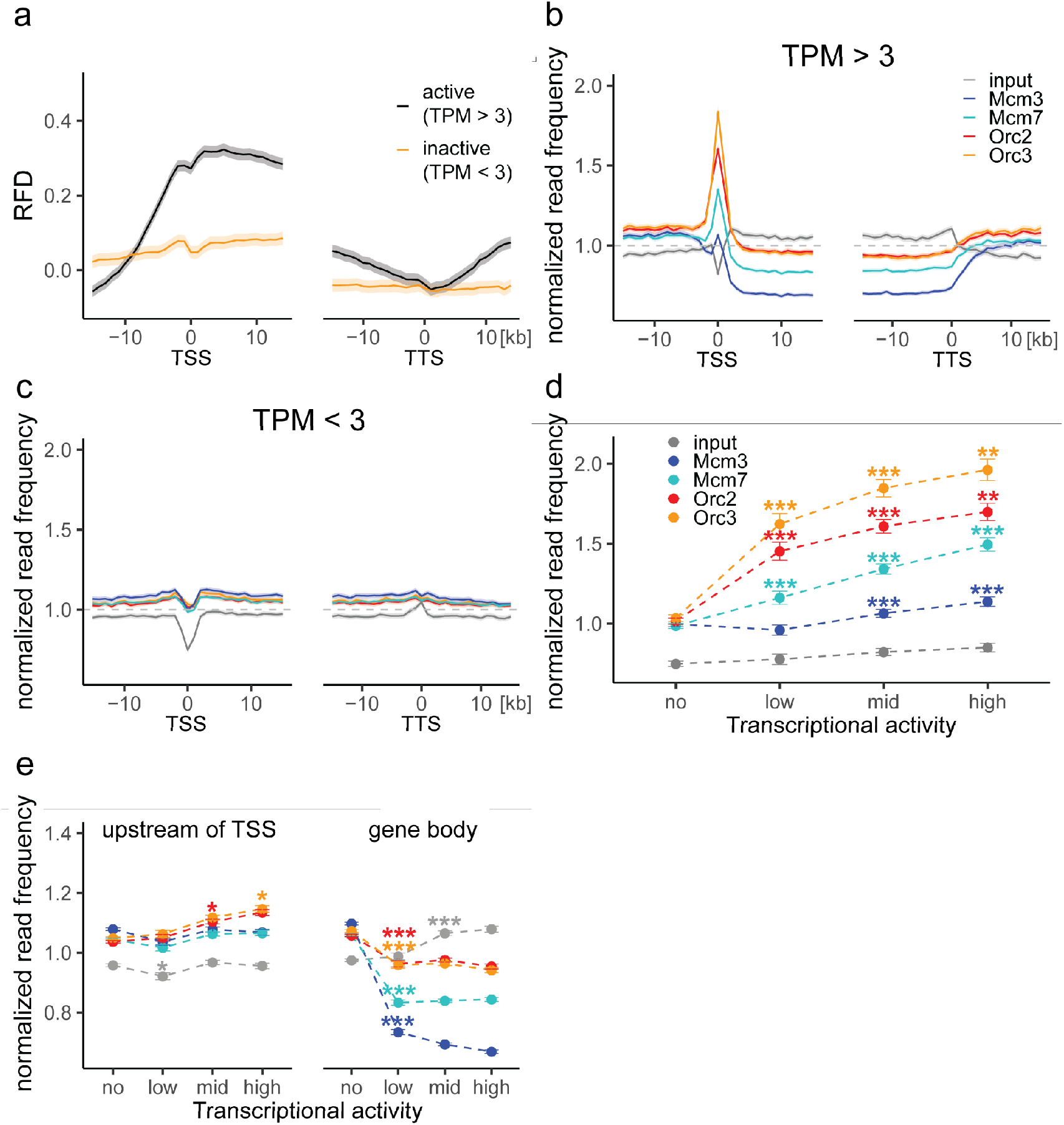
ORC and MCM profiles at genes without input normalization. a) RFD around TSSs or TTSs of active genes (black) or inactive genes (yellow). Distances from TSSs or TTSs are indicated in kb. RFD means are shown ± 2 x SEM (lighter shadows). b - c) Normalized ORC/MCM/input read frequencies without input division around TSSs or TTSs for b) active genes (TPM > 3) and c) inactive genes (TPM < 3). Only genes larger than 30 kb without any adjacent gene within 15 kb were considered. Distances from TSSs or TTSs are indicated in kb. Means of normalized read frequencies are shown ± 2 x SEM (lighter shadows). The dashed grey horizontal line indicates read frequency 1.0 for reference. d) Normalized ORC/MCM/input read frequencies at TSSs dependent on transcriptional activity (± 2 x SEM). e) Normalized ORC/MCM/input read frequencies upstream of TSSs and in the gene body dependent on transcriptional activity (± 2 x SEM; TSSs ± 3 kb removed from analysis). Transcriptional activity was classified as: no (TPM < 3), low (TPM 3-10), mid (TPM 10-40), high (TPM > 40). Statistics were performed by one-way ANOVA followed by Tukey’s post-hoc test. P-values are indicated always comparing to the previous transcriptional level. * p < 0.05, ** p < 0.01, *** p < 0.001.

**Supplementary Figure 7:**
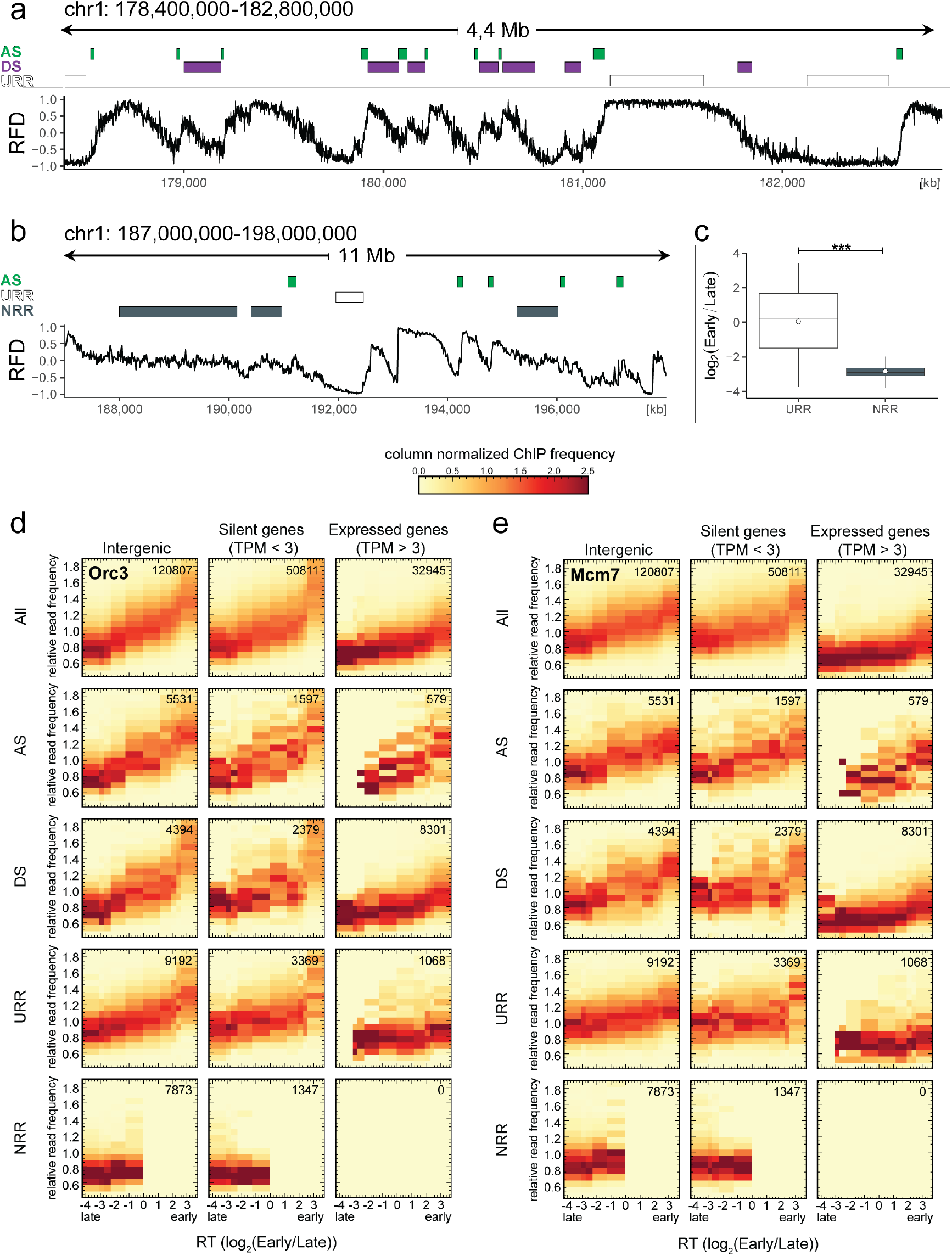
ORC/MCM levels are correlated with RT and transcriptional activity but otherwise homogeneously distributed along the genome and uncorrelated to RFD patterns. a) Same RFD profile example as Fig. 2a (chr1: 178,400,000 – 182,800,000, covering 4 Mb) with indication of AS (green), DS (purple), and URR (white boxes) positions. b) RFD profile example on chr1: 187,000,000 – 198,000,000, covering 11 MB, with ASs (green), URRs (white boxes), and NRRs (dark gray) indicated. c) Replication timing ratio log2(Early/Late) was assigned to URRs and NRRs and represented as boxplot (mean (circle), the median (thick line), the 1st and 3rd quartile (box), the 1st and 9th decile (whiskers), without representing outliers). Statistics were performed using one-sided t-test. *** p < 0.001. d-e) 3×5 panel of 2D histograms representing Orc3 (d) and Mcm7 (e) ChIP frequency vs. RT (average log2(Early/Late) over 100 kb binned according to the decile of RT distribution). The analysis was performed in 10 kb windows. ChIP relative read frequencies are normalized by column and represent the probability density function of ChIP frequency at a given replication timing. The color legend is indicated on top. The columns of each panel represent only windows present in intergenic regions (left column), silent genes (TPM < 3, middle column), and expressed genes (TPM > 3, right column). TSSs and TTSs proximal regions were not considered (see Material and Methods). The rows show either all bins (top row), AS bins (predominant replication initiation, second row), DS bins (descending segment, predominant replication termination, third row), URR bins (unidirectional replication, no initiation, no termination, fourth row) and NRR bins (null RFD regions, spatially random initiation and termination, bottom row). The number of bins per panel is indicated in each panel. See Fig. 4 for equivalent Orc2 and Mcm3 data. Refer to Suppl. Fig. 8a for statistical significances.

**Supplementary Figure 8:**
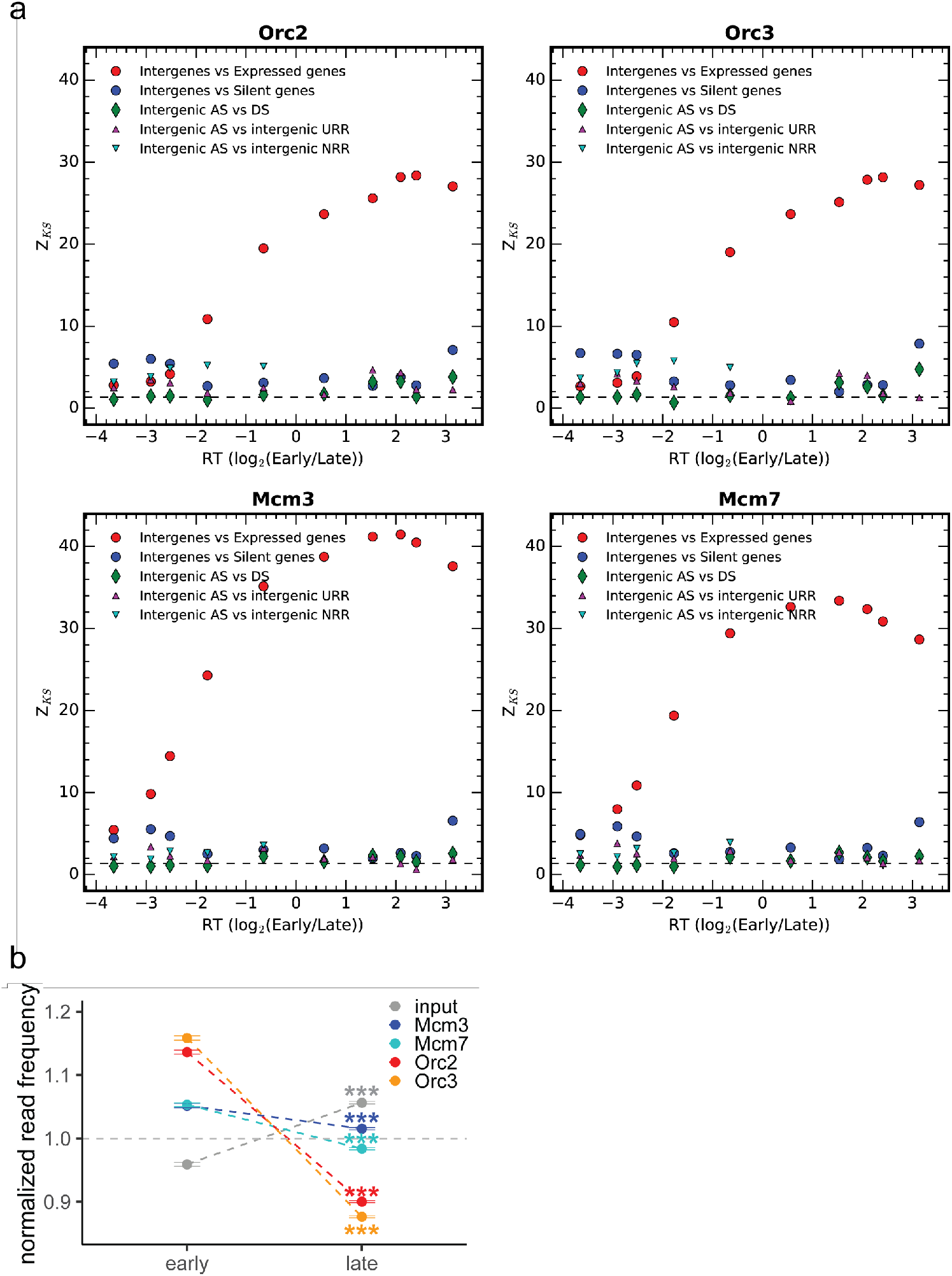
a) Kolmogorov–Smirnov statistics between the ORC/MCM relative read frequency distributions in each replication timing bin (shown in Fig. 4 and Suppl. Fig. 7d and 7e) in intergenic vs expressed gene regions (red circles), in intergenic versus silent gene regions (blue circles) and between intergenic regions in AS versus DS (green diamonds), in AS versus URR (magenta triangles pointing up) and in AS versus NRR (cyan triangles pointing down). Z_KS is normalized for sample size. The horizontal dashed lines correspond to p-value = 5%. b) Normalized ORC/MCM/input read frequencies without input division (± 2 x SEM) in early or late RTDs. Early RTDs were defined as log2(Early/Late) > 1.6; late RTDs < -2.0. The analysis was performed in 10 kb bins. Any gene ± 10 kb was removed from the analysis. Statistics were performed using one-sided t-test. *** p < 0.001.

**Supplementary Figure 9:**
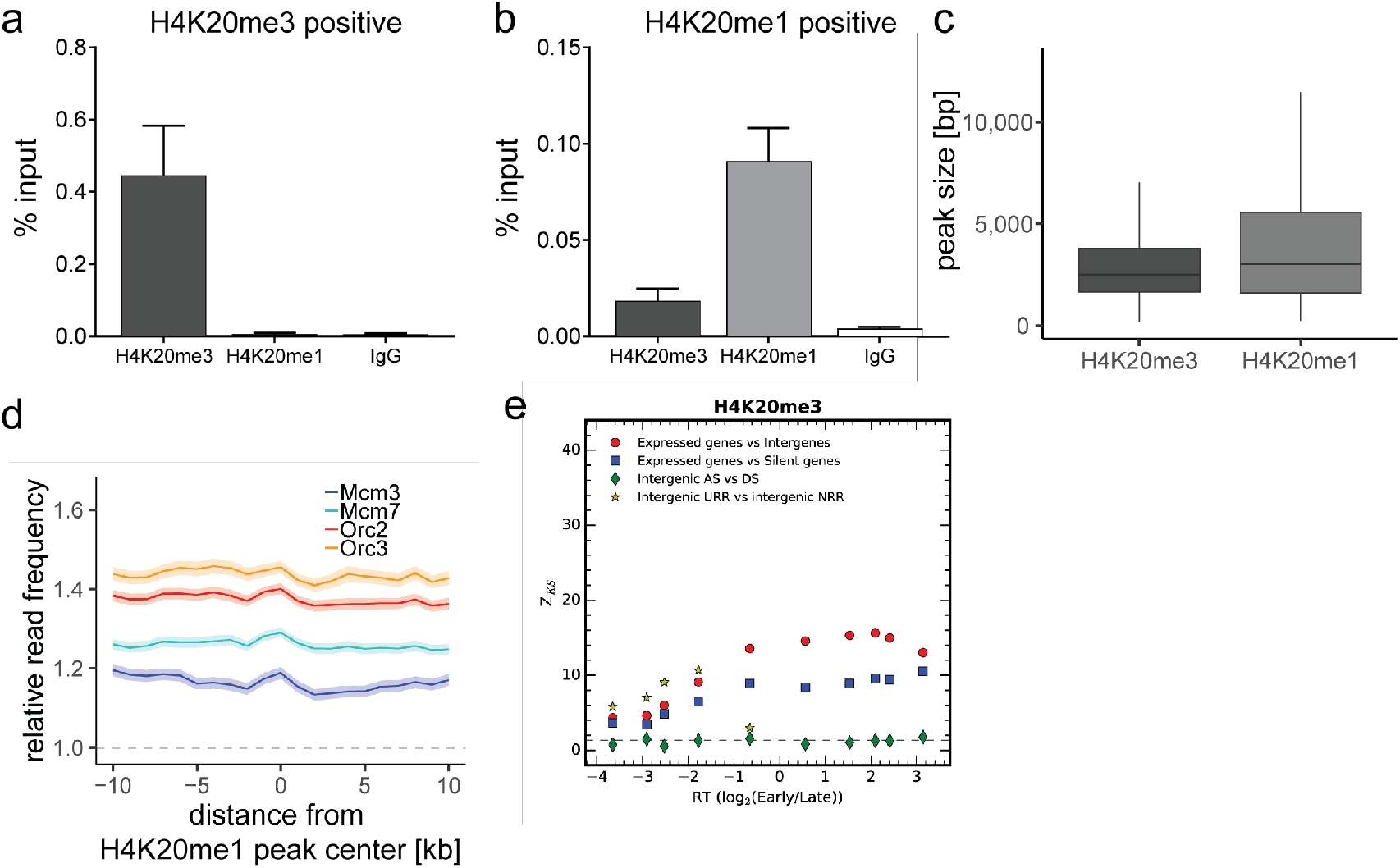
ORC/MCM is enriched in late replicating, H4K20me3-high non-genic AS and NRR windows. a)-b) qPCR validation of H4K20me3 and H4K20me1 enrichment after ChIP at a) an H4K20me3 positive locus and b) an H4K20me1 positive locus. Representation in % input. Isotype IgG antibodies were used as control. c) Boxplot of H4K20me3 and H4K20me1 peak size (in bp) distribution. d) Average ORC/MCM relative read frequencies after input normalization at H4K20me1 peaks (> 1 kb). e) Kolmogorov–Smirnov statistics between the H4K20me3 relative read frequency distributions in each replication timing bin (shown in Fig. 6a): expressed gene versus intergenic (red circles) and silent gene (blue squares) regions as well as between intergenic regions in AS versus DS (green diamonds) and in URR versus NRR (yellow stars). Z_KS is normalized for sample size. The horizontal dashed lines correspond to p-value = 5%.

**Supplementary Figure 10:**
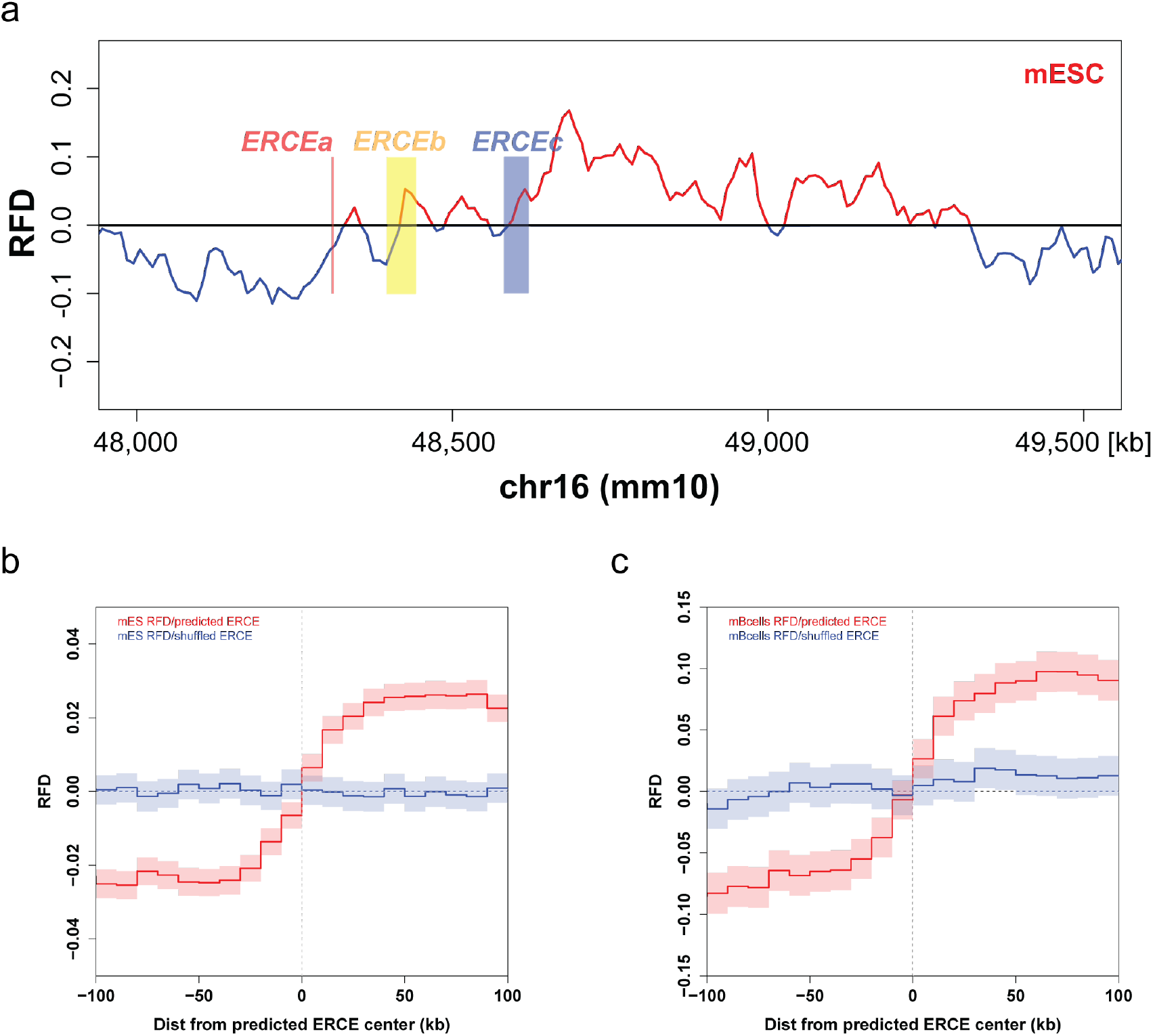
Early replication control elements (ERCEs) correlate with replication initiation. a) mESC OK-seq RFD profile of the mouse Dppa2/4 locus (chr16: 48,000,000 – 49,500,000) with indicated ERCEs (ERCEa, ERCEb, ERCEc). ERCEa and ERCEc are located within ascending RFD segments (AS) in mESCs. ERCEb encompasses an entire AS. b) Mean mESC OK-seq RFD profile around the 1,835 mESC ERCEs predicted by (Sima et al., 2019) (red), and randomly shuffled ERCEs (blue). c) Mean mouse primary B-cell OK-seq RFD profile around the same mESC ERCE set (red) and randomly shuffled ERCEs (blue). mESC OK-seq data was obtained from (Petryk et al., 2018); mouse primary B cell OK-seq data was computed from Tubbs et al., 2018; mESC ERCEs were predicted by (Sima et al., 2019).

**Supplementary Table 1:** Proportion of genes significantly depleted from ORC/MCM. A total of 1,941 genes met the criteria of transcriptional activity (TPM > 3), gene size larger 30 kb and no adjacent genes within 15 kb. We calculated the proportion of genes where the mean relative read frequency within the gene was significantly (p < 0.05) reduced compared to the upstream region (excluding TSS +/− 3 kb).

| | Total genes | $p < 0.05$ | % |
| --- | --- | --- | --- |
| Orc2 | 1,941 | 865 | 44.6 |
| Orc3 |  | 850 | 43.8 |
| Mcm3 |  | 1,460 | 75.2 |
| Mcm7 |  | 1,131 | 58.3 |

**Supplementary Table 2:** Ratio of ChIP mean relative read frequencies in early vs. late RTDs. Calculated in 10 kb bins. All annotated genic regions were removed ± 10 kb.

|  | Mean relative read frequency ratio<br>(early/late) |
| --- | --- |
| Orc2 | 1.40 |
| Orc3 | 1.47 |
| Mcm3 | 1.15 |
| Mcm7 | 1.19 |

**Supplementary Table 3:**
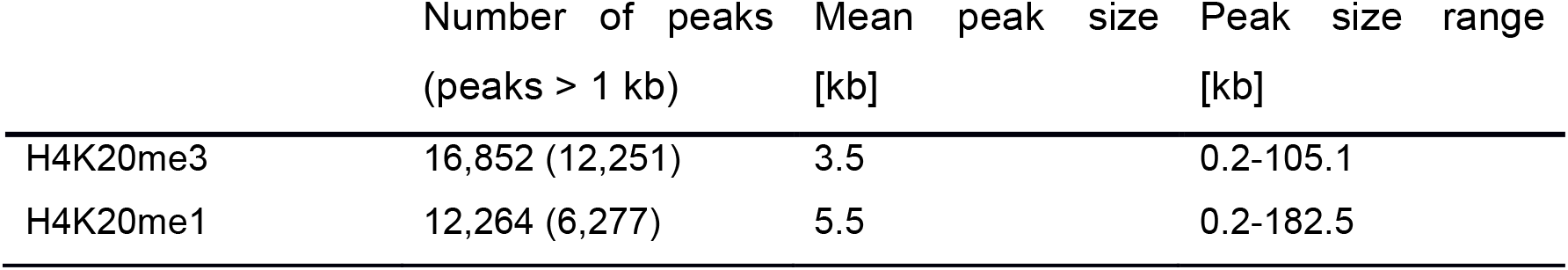
Characterization of H4K20me3 and H4K20me1 peaks determined by MACS2 broad peak calling.

